# lncRNA H19/Let7b/EZH2 axis regulates somatic cell senescence

**DOI:** 10.1101/2022.07.07.499142

**Authors:** Manali Potnis, Justin Do, Olivia El Naggar, Eishi Noguchi, Christian Sell

**Affiliations:** Department of Biochemistry & Molecular Biology, Drexel University College of Medicine, Philadelphia, PA 19102

## Abstract

Long non-coding RNAs (lncRNAs) regulate diverse cellular processes and are associated with many age-associated diseases. However, the function of lncRNAs in cellular senescence remains largely unknown. Here we characterize the role of lncRNA H19 in senescence. We show that H19 levels decline as cells undergo senescence, and depletion of H19 results in premature senescence. We find that repression of H19 is triggered by the loss of CTCF and prolonged activation of p53 as part of the senescence pathway. Mechanistically, the loss of H19 drives senescence via increased let7b mediated targeting of EZH2. We further demonstrate that H19 is required for senescence inhibition by the mTOR inhibitor rapamycin, where it maintains lncRNA H19 levels throughout the cellular lifespan and thus prevents the reduction of EZH2 that would otherwise lead to cellular senescence. Therefore, lncRNA H19 is crucial in maintaining the balance between sustained cell growth and the onset of senescence.

## Introduction

Cellular senescence is a programmed cell fate characterized by a stable cell-cycle arrest. Initially described in primary cells after long-term cell culture, it was attributed to telomere attrition (1, 2). However, it is now clear that the telomere paradigm is an oversimplification and that cellular senescence is a response to multiple types of stress such as DNA double-strand breaks, metabolic imbalance, and oncogene activation (3-5). Accumulating evidence indicates a potential causal role for senescence in organismal aging (6, 7). Senescent cells increase with age in multiple tissues and are associated with age-related disorders such as Alzheimer’s disease, cardiovascular disease, and arthritic changes (6), while direct targeting of senescent cells provides late-life benefits. For example, the depletion of senescent cells relieves symptoms in mouse models of age-related diseases (8-10), suggesting that targeting cellular senescence is a viable approach to improving organismal aging and alleviating age-related disorders such as cardiomyopathy and cognitive decline. Although senescence can be deleterious, there is evidence that senescence may also play beneficial roles in physiologic processes. For example, senescence occurs during development (11, 12), and senescent cells appear to contribute to proper wound healing (13). Interestingly, an evaluation of senescence in mammalian species suggests a positive correlation between the propensity of mammalian cells to undergo senescence and longevity (14), suggesting a positive role for senescence in longevity. We have demonstrated that the mTOR inhibitor, rapamycin, can delay cellular senescence and modify the senescent phenotype in human cells (15-17), reduce pro-inflammatory cytokine production, and improve metabolic balance (18, 19) as well as alter chromatin-modifying enzymes (20).

The senescence program requires large-scale spatial rearrangement of chromatin, This rearrangement varies depending on the cell type and the specific stress used to induce senescence (21), resulting in condensation of some areas of the genome (22) but reduced compaction in other regions (21, 23, 24). As additional modifiers of chromatin structure are identified, novel aspects of regulating chromatin architecture in senescence are emerging. Long non-coding RNAs (lncRNAs) have multiple functions in the nucleus related to chromatin remodeling (25), including the recruitment of chromatin-modifying complexes such as PRC2 (26, 27), and there is evidence for the involvement of a subset of non-coding RNAs in senescence (28, 29). For example, the lncRNA MIR31HG regulates the Ink4A locus during oncogene-induced senescence (30), and HOTAIR activates cellular senescence through p53-p21^Cip1/Waf1^ signaling of the DDR in chemotherapy-resistant ovarian cancer cells (31). The lncRNA GUARDIN is required for genomic stability, and targeting of GUARDIN induces senescence (32). lncRNAs appear to also play a direct role in the senescence program, and elucidating their function is essential to our understanding of both senescence and aging.

H19 is a highly conserved, maternally expressed imprinted gene and encodes a 2.3 kb long non-coding RNA (lncRNA). It is located immediately downstream of the neighboring gene IGF2. Functionally, H19, in conjunction with the neighboring gene, IGF2, is critical in controlling embryonic growth (33, 34). Loss of H19 results in fetal overgrowth associated with Beckwith-Weidemann syndrome(35), while elevated H19 occurs in human cancers (36, 37). In a recent study, H19 was shown to be downregulated in aged cardiac endothelial cells (38), and while most adult tissues experience a substantial decline in H19 levels with age (38-41), the cause and consequence of the reduced H19 levels are unclear.

Here we show that by preventing let7b-mediated degradation of EZH2, lncRNA H19 maintains somatic cell proliferation and provides a buffer against senescence. Subsequently, H19 depletion, induced by loss of CTCF or p53 activation, leads to senescence via the H19/let7/EZH2 axis, suggesting that H19 acts as a critical node that maintains the balance between sustained cell growth and the onset of senescence. Importantly, H19 is also required for the senescence delaying effects of rapamycin in somatic cells.

## Results

### Suppression of H19 induces premature senescence

To characterize the role of H19 in the cellular senescence of somatic cells, we examined H19 expression during replicative senescence of human cardiac fibroblasts. We observed a significant decrease in lncRNA H19 levels in late passage cells relative to early passage cells **(Figure 1a)**. Remarkably, treatment with rapamycin, which we have previously shown to prevent the transition of cells into a senescent state (42), maintained H19 levels in late passage cells, and discontinuing the treatment led to a reduction in H19 levels **(Figure 1a, Figure 1-figure supplement 1a)**. To confirm whether these changes are reflected in aged tissues, we analyzed skin samples from old mice treated with or without topical rapamycin treatment. H19 expression was lower in aged (20 months) mice relative to young (3 months) and was significantly elevated in aged skin treated with rapamycin **(Figure 1b)**. The results indicate that H19 levels are reduced during aging in vitro and in vivo and that H19 levels are maintained by rapamycin treatment.

**Fig 1:**
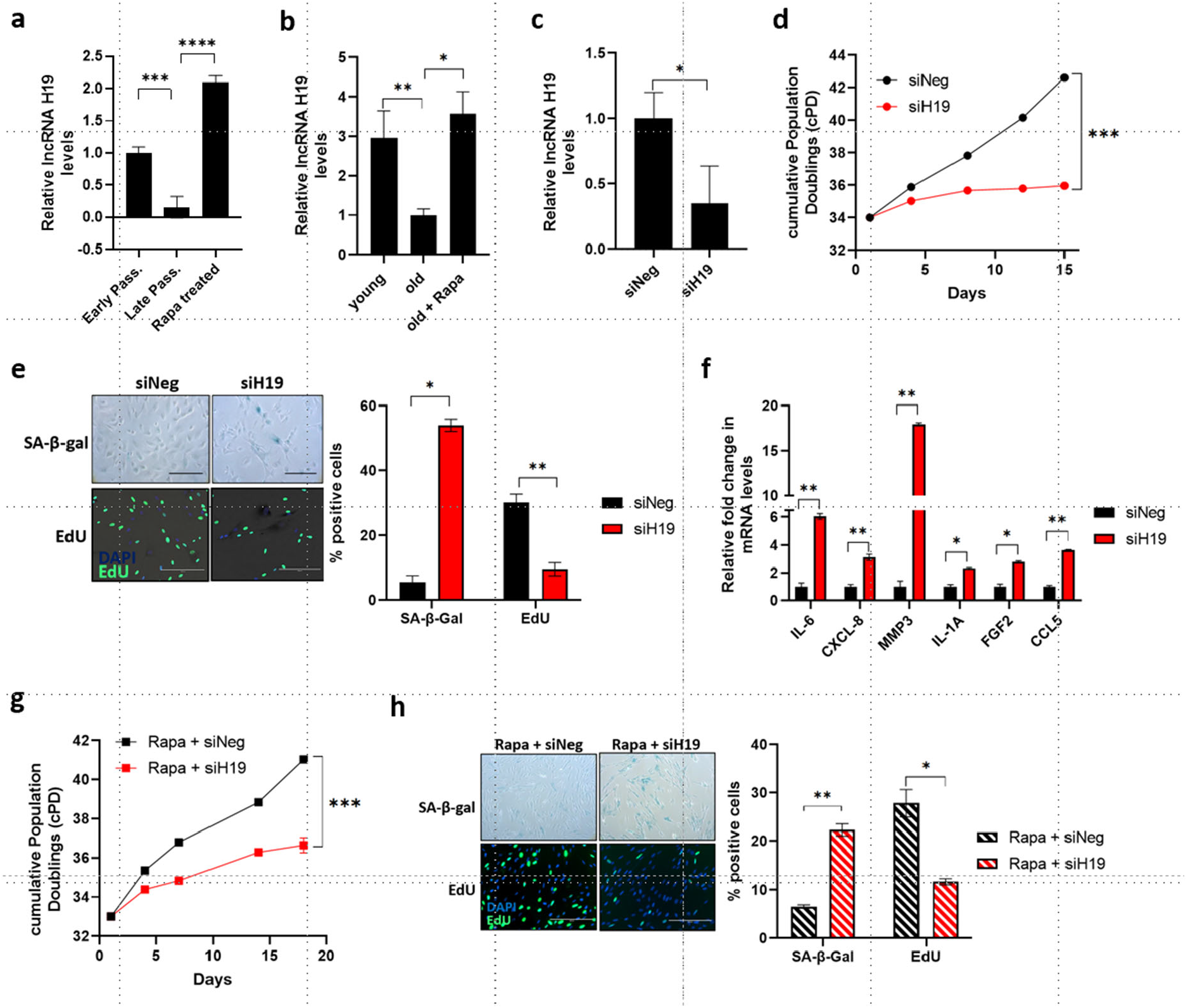
H19 expression is reduced during senescence but maintained with rapamycin treatment. **a** lncRNA H19 steady-state levels were determined by RT-qPCR in human cardiac fibroblasts (HCF) at early passage, late passage, and late passage treated with rapamycin (Mean±SEM, **p < 0.01 by two-tailed unpaired Student’s *t*-test, n = 3). **b** lncRNA H19 levels were determined by RT-qPCR in the skin of young, old, and old mice treated with topical rapamycin (Mean±SEM, *p < 0.05 by two-tailed unpaired Student’s *t*-test, **p < 0.01, n = 3). **c** Early passage HCF cells were transfected with siRNA targeting H19 (siH19) and negative control (siNeg), and lncRNA H19 levels were determined 4 days after transfection (Mean±SEM, *p < 0.05 by two-tailed unpaired Student’s *t*-test, n = 3). **d** H19 was knocked down in early passage HCF cells as in c, and growth was assessed by cumulative population doublings (cPD) (***p < 0.001 by two-tailed unpaired Student’s *t*-test, n = 3). **e** SA-β-gal activity and DNA synthesis were analyzed in H19 knockdown cells by SA-β-gal and EdU staining, respectively. Left: Images of SA-β-gal and EdU stained control (siNeg) and H19KD (siH19) HCF cells are shown. The scale bar represents 200 µm. Right: Quantified data is represented as a percentage of positive cells (Mean±SEM, *p < 0.05 and **p < 0.01 by two-tailed unpaired Student’s *t*-test, n = 3). **f** Senescence-associated secretory phenotype (SASP) in H19 targeted cells was assessed by RT-qPCR (Mean±SEM, *p < 0.05 and **p < 0.01 by two-tailed unpaired Student’s *t*-test, n = 3). **g** H19 was knocked down in early passage cells maintained in rapamycin-containing media, and growth was assessed by cumulative population doublings (cPD) (***p < 0.001 by two-tailed unpaired Student’s *t*-test, n = 3). **h** SA-β-gal activity and DNA synthesis were analyzed in H19 knockdown cells treated with rapamycin by SA-β-gal and EdU staining, respectively. Left: Images of SA-β-gal and EdU stained control (Rapa + siNeg) and H19KD (Rapa + siH19) cells are shown (Mean±SEM, *p < 0.05 and **p < 0.01 by two-tailed unpaired Student’s *t*-test, n = 3). Right: Quantified data is represented as a percentage of positive cells. Source data are provided as a Source Data file.

We next sought to verify whether targeting H19 would trigger the senescence program. Consistent with the role of H19 in senescence, depleting H19 using siRNAs **(Figure 1c; Figure 1-figure supplement 1b)** decreased the life span of cells **(Figure 1d; Figure 1-figure supplement 1c)** and triggered cellular senescence characterized by an increase in SA-β-gal activity, and suppressed proliferation indicated by decreased incorporation of EdU **(Figure 1e; Figure 1-figure supplement 1d)**. In addition, H19 downregulation increased the expression of SASP-associated genes **(Figure 1f)**. Although rapamycin restored H19 expression in late passage cells, rapamycin treatment was not enough to rescue the onset of senescence following direct targeting of H19 **(Figure 1g and h; Figure 1-figure supplement 1e and f)**, suggesting that H19 is essential for the inhibitory effect of rapamycin on cellular senescence. Interestingly, overexpressing H19 triggered senescence as demonstrated by decreased proliferation and increased SA-β-gal activity **(Figure 1-figure supplement 1g)**. Taken together, these findings suggest that H19 levels are maintained within a critical range to maintain cell function. These results are comparable to studies examining the role of Lamin B1 during senescence, where a reduction in Lamin B1 levels accompanies senescence, and either depletion or overexpression of Lamin B1 induces senescence (43).

### Loss of CTCF expression during senescence results in decreased H19 expression

To explore the molecular mechanism underlying the regulation of H19 in cellular senescence, we sought to identify regulators of H19. Binding of the CCCTC-binding factor (CTCF) to the H19 regulatory imprinted control region (H19 ICR) is thought to regulate allelic H19 and IGF2 gene expression where the unmethylated maternal H19 ICR bound to CTCF allows for H19 gene expression and hypermethylated paternal H19 ICR allows IGF2 gene expression (44-47). Downregulation of CTCF expression has been demonstrated to accompany the onset of senescence (48-50). Consistent with these published results, CTCF mRNA and protein levels decreased in the late passage cells **(Figure 2a and b)**, and CTCF knockdown in early passage cells induced premature senescence characterized by increased SA-β-gal staining and reduction in DNA synthesis **(Figure 2-figure supplement 2a)**. In contrast, treatment with rapamycin prevented CTCF loss, which is consistent with the effect of rapamycin maintaining H19 levels **(Figure 2a and b)**. Furthermore, the regulatory link between CTCF and H19 is supported by decreased H19 expression in CTCF-targeted cells **(Figure 2c)**. These results confirm that reduction in CTCF is at least partly responsible for the downregulation of H19 expression as part of the senescence pathway.

**Fig 2:**
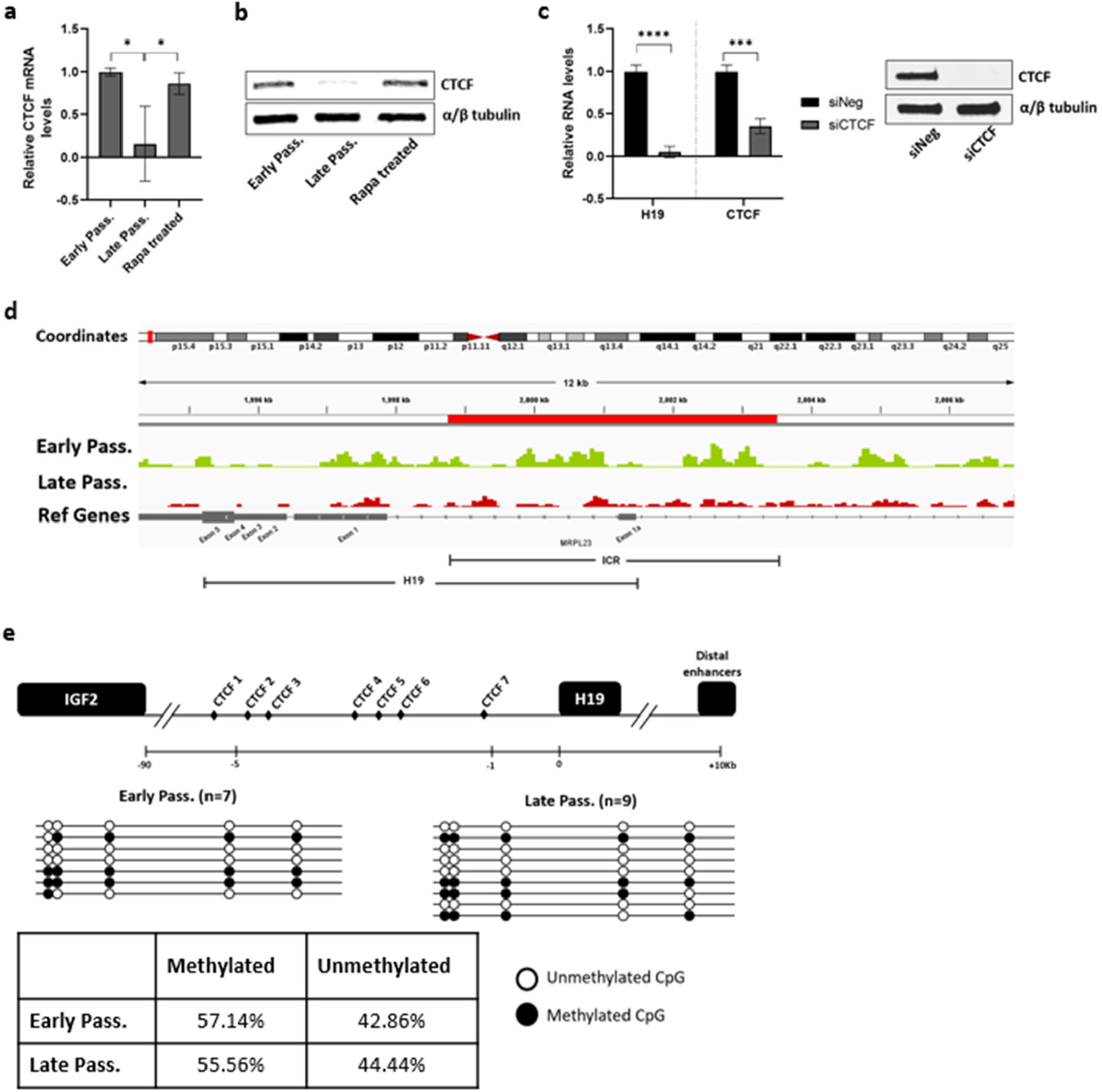
Loss of CTCF leads to a reduction in H19 expression. **a** CTCF mRNA expression was determined by RT-qPCR in early passage andlate passage HCFs, or late passage HCFs treated with rapamycin (Mean±SEM, *p < 0.05 by two-tailed unpaired Student’s *t*-test, n = 3). **b** CTCF protein expression was analyzed by western blot in early, late, and late passage HCFs treated with rapamycin. α/β tubulin serves as the loading control. **c** Early passage HCF cells were transfected with siRNA targeting CTCF (siCTCF) and negative control (siNeg). Right: lncRNA H19 and CTCF mRNA levels were determined 7 days after transfection (Mean±SEM, ***p < 0.001 and ****p < 0.0001 by two-tailed unpaired Student’s *t*-test, n = 3). Left: Western blot images of CTCF protein in the early passage HCF cells transfected with siCTCF. α/β tubulin serves as the loading control. **d** Representative tracks from CTCF CUT&Tag-sequencing for H19. Tracks show CTCF binding at the H19 gene and H19-ICR for early and late passage HCF cells in green and red, respectively. The Refseq gene track is displayed in grey. The H19-ICR is shown in red. **e** Structure for the human H19-ICR is shown. Putative CTCF and other binding sites are indicated. The methylation status of CTCF binding site 5 was obtained by bisulfite sequencing followed by PCR amplification and cloning of individual PCR products. Each line represents an individual clone with indicated early and late passage status. Methylated CpGs are shown as black circles and unmethylated ones as white circles. The percentage of methylated and unmethylated clones is shown in the adjacent table. Source data are provided as a Source Data file.

Recent studies have described a stress-dependent downregulation of CTCF through proteasomal degradation of CTCF protein in endothelial cells (51). To validate whether increased proteasomal degradation contributes to the reduction of CTCF protein observed during senescence, we treated late passage cells with the proteasomal inhibitor MG132. However, no change in CTCF protein levels was noted in the late passage cells treated with MG132 despite a general increase in polyubiquitinated proteins, suggesting that stress-dependent proteasomal clearance of CTCF does not contribute to the reduction in CTCF protein during senescence of HCFs **(Figure 2-figure supplement 2b)**.

The loss of binding of CTCF at the IGF2-H19 locus is critical in regulating the biallelic expression of IGF2 and H19 in mice (44). Moreover, loss of CTCF binding at the H19-ICR has been demonstrated to contribute to increased IGF2 expression in senescent human epithelial cells (48). To evaluate if a reduction in CTCF levels during senescence results in decreased binding of CTCF at the H19-ICR in our system, we subjected proliferating early passage and late passage cells to CUT & Tag-sequencing. A comparison of CTCF binding profiles at the H19-ICR in early and late passage HCF cells shows a relative reduction in CTCF binding at the H19-ICR in late passage cells **(Figure 2d)**. Additionally, in humans, silencing of H19 expression is associated with de novo methylation or hypermethylation of H19 ICR, as demonstrated in Wilms’ tumors (52-54) and colorectal cancers (55). To investigate if hypermethylation at the maternal H19 ICR contributes to the loss of H19 expression during senescence, we performed a differential methylation analysis of the sixth CTCF binding site in the H19 ICR, which has been identified to undergo allele-specific differential methylation and loss of imprinting in cancers (52, 55-57). Allelic methylation at the ICR predicts 50% methylation in the normal cells, and an increase from 50% suggests hypermethylation of the unmethylated maternal ICR. However, following bisulfite treatment and Sanger sequence analysis of the early and late passage cells, no methylation changes were noted at the sixth CTCF binding site in the unmethylated allele **(Figure 2e)** indicating differential methylation of the H19 ICR does not underlie the loss of H19 in senescent cells.

Senescence is associated with increased IGF2 gene expression (48) **(Figure 2-figure supplement 2c)**. To examine if IGF2 drives senescence in H19 depleted cells, we examined IGF2 levels in H19 targeted cells which previously demonstrated activation of the senescence program. Although depleting CTCF increased IGF2 mRNA levels **(Figure 2-figure supplement 2d)**, targeting H19 did not affect IGF2 mRNA levels suggesting that depletion of H19 induced premature senescence independent of IGF2 **(Figure 2-figure supplement 2e)**.

### Prolonged activation of p53 results in decreased H19 expression

Given the stress-responsive nature of CTCF (51), we then evaluated whether loss of CTCF or H19 occurs during the DNA damage response (DDR). Although we observed a reduction in H19 levels in response to acute treatment with genotoxic agent Neocarzinostatin (NCS) and oxidative stressor hydrogen peroxide (H2O2) **(Figure 3a)**, there was no change in CTCF protein levels **(Figure 3b)**, implying that other mechanisms regulate H19 in response to acute stress.

**Fig 3:**
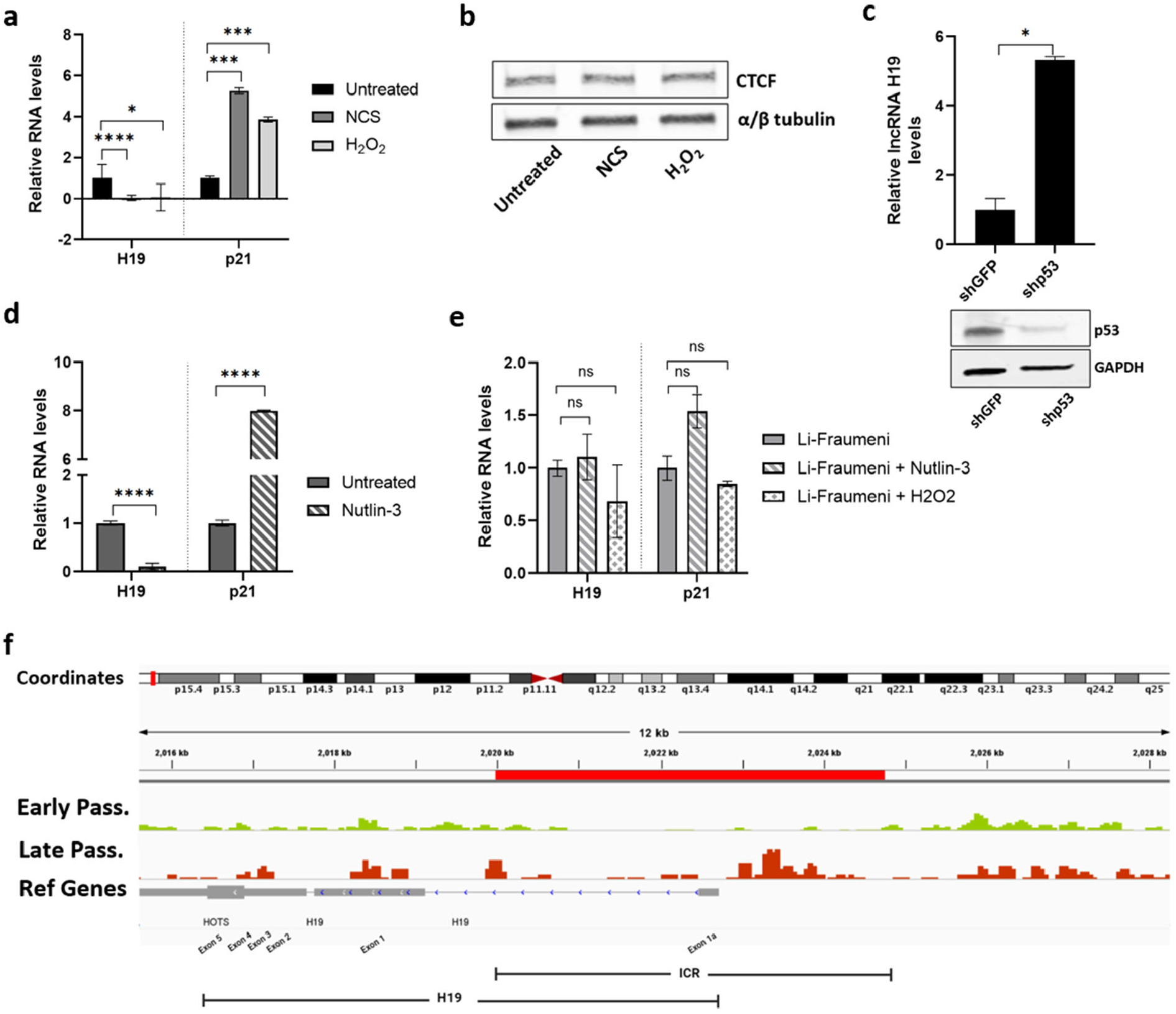
p53 activation results in decreased H19 expression. **a** Early passage HCF cells were treated with genotoxin neocarzinostatin (0.2μM) and hydrogen peroxide (200μM), and lncRNA H19 and p21^Cip1/Waf1^ mRNA levels were determined 24 hrs after treatment (Mean±SEM, **p < 0.05, **p < 0.001, and ****p < 0.0001 by two-tailed unpaired Student’s *t*-test, n = 3). **b** CTCF protein expression was analyzed by western blot in cells treated with Neocarzinostatin and hydrogen peroxide (24hr post-treatment). α/β tubulin serves as the loading control. **c** Early passage HCF cultures were infected with control (shGFP) and p53 knockdown (shp53). Top: lncRNA H19 levels were determined in HCF cells infected with shGFP and shp53 (Mean±SEM, *p < 0.05 by two-tailed unpaired Student’s *t*-test, n = 3). Bottom: Western blot images of p53 protein in HCF cell infected with shGFP and shp53. GAPDH serves as the loading control. **d** Early passage HCF cells were treated with MDM2 inhibitor Nutlin-3, and lncRNA H19 and p21^Cip1/Waf1^ mRNA levels were determined 72 hrs after treatment (Mean±SEM, ****p < 0.0001 by two-tailed unpaired Student’s *t*-test, n = 3). **e** lncRNA H19 and p21^Cip1/Waf1^ mRNA levels were determined by RT-qPCR in the Li-Fraumeni cells treated with Nutlin-3 and hydrogen peroxide (Mean±SEM, p > 0.05 by two-tailed unpaired Student’s *t*-test, n = 3). Source data are provided as a Source Data file. **f** Representative tracks from p53 CUT&Tag-sequencing for H19. Tracks show p53 binding at the H19 gene and H19-ICR for early and late passage HCF cells in green and red, respectively. The Refseq gene track is displayed in grey. The H19-ICR is shown in red.

Activation of p53 is crucial for establishing senescence as part of DDR. Importantly, p53-dependent repression of the H19 promoter has been demonstrated in stem cells and cancers (58, 59). To verify whether p53 regulates H19 in normal somatic cells, we analyzed H19 levels in p53-depleted cells and p53-activated cells. Downregulation of p53 resulted in increased H19 levels **(Figure 3c)**, whereas Nutlin-3-mediated p53 activation led to a decline in H19 levels **(Figure 3d)**. Consistent with these findings, Li Fraumeni cells harboring p53 LOF mutation (TP53 R175h mutation) displayed unchanged levels of H19 despite treatment with Nutlin-3 and oxidative agents **(Figure 3e)**. Further, despite the lack of conventional TATA-box and p53 consensus binding sites in the H19 promoter, wild-type p53 has been shown to downregulate the activity of the H19 gene promoter (58). To evaluate if increased p53 during senescence is reflected in changes in binding of p53 at the H19 promoter, we subjected proliferating early passage and late passage cells to CUT & Tag-sequencing. A comparison of p53 binding profiles at the H19 in early and late passage HCF cells shows a relative increase in p53 binding at the H19 promoter **(Figure 3f)** and downstream gene target p21Cip1/Waf1 **(Figure 3-figure supplement 3a)** in late passage cells. Together these results confirm that activation of p53 is responsible for the downregulation of H19 as part of DNA damage response.

### Decreased H19 induces senescence via the let7b/EZH2 axis

Next, we sought to dissect the potential downstream mechanisms which trigger the senescence state in H19 depleted cells. Given the mounting evidence suggesting the role of lncRNA H19 as a competing endogenous RNA (ceRNA) or miRNA sponge (60-62), we speculated that H19 might mediate the senescence program by regulating miRNA availability. To determine which miRNAs are directly regulated by lncRNA H19 during senescence, we evaluated miRNA expression profiles in control, and H19 targeted cells **(Figure 4a)**. Among the top miRNAs upregulated in H19 depleted cells were members of the let7 family; specifically, let7b expression was significantly upregulated **(Figure 4b, Figure 4-figure supplement 4a)**. This is supported by the evidence of direct interaction between lncRNA H19 and let7b via complementary binding sites (60, 63) **(Figure 4-figure supplement 4b)**. In addition, an up-regulation of let7b is observed in aged human stem cells and primary muscle cells (64-67). Based on these data, we examined let7b levels in HCFs during senescence and found that let7b levels increased in the late passage cells and consequently decreased with rapamycin **(Figure 4c)**. Additionally, increased let-7b levels using synthetic let-7b-5p mimics induced premature senescence in young proliferating HCFs, which was characterized by cessation of growth **(Figure 4d)**. Further, let-7b-5p mimics decreased proliferation, increased SA-β-gal activity **(Figure 4e)**, and elevated senescence-associated secretory phenotype (SASP) mRNA levels **(Figure 4f)**. Taken together, these data confirmed increased levels of let-7b post depletion of lncRNA H19 and support the concept that let-7b acts as a downstream target of lncRNA H19 during senescence.

**Fig 4:**
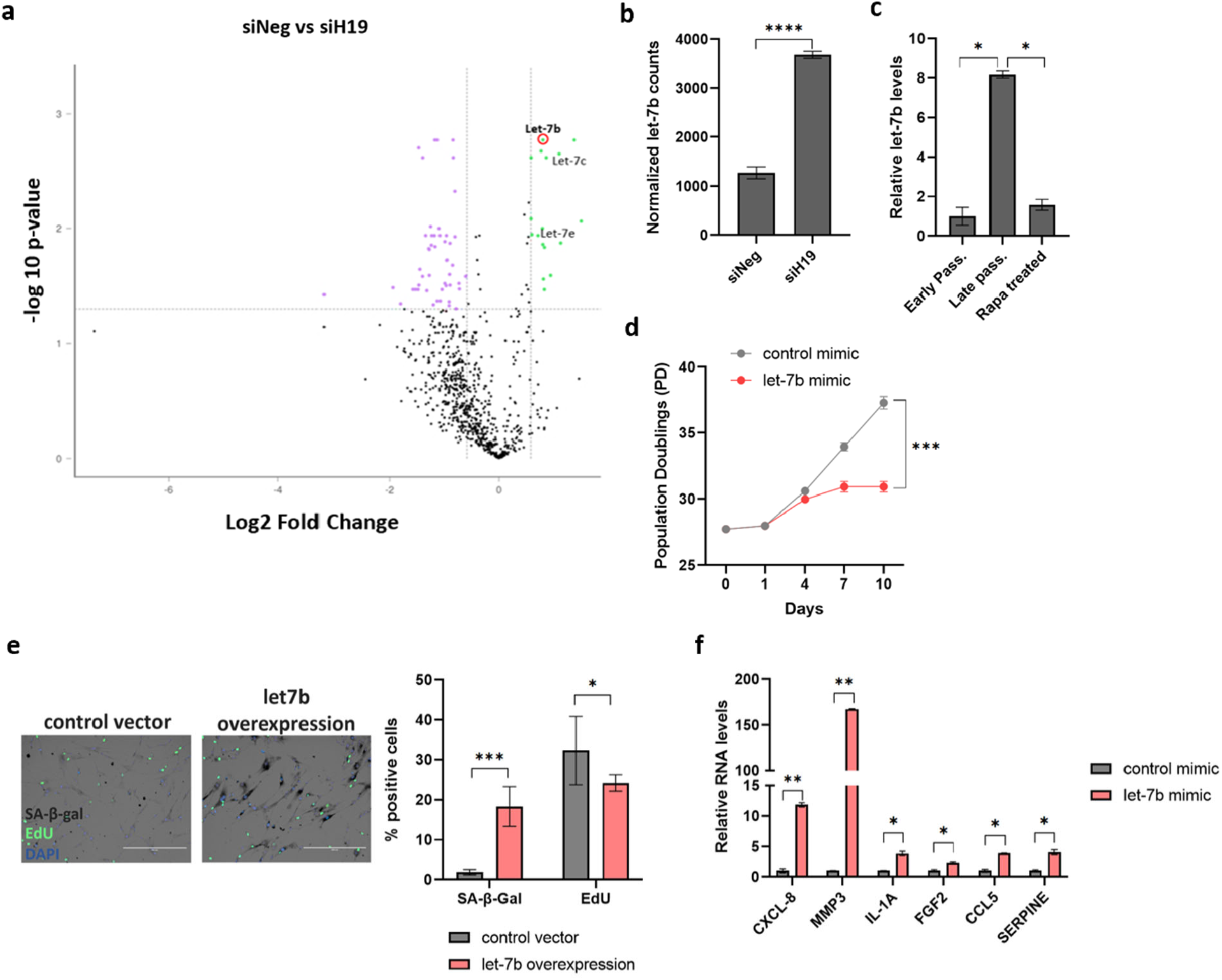
let-7b/EZH2 axis in H19 depletion and senescence. **a** Volcano plot based on a Nanostring analysis of the expression of ∼800 miRNAs upon H19 targeting in HCF cells vs. control. **b** Normalized counts of let-7b from Nanostring analysis in siH19 and siNeg targeted cells (Mean±SEM, ****p < 0.0001 by two-tailed unpaired Student’s t-test, n = 3). **c** Let-7b expression was determined by RT-qPCR in the early passage, late passage, and late passage human cardiac fibroblasts (HCF) treated with rapamycin (Mean±SEM, *p < 0.05 by two-tailed unpaired Student’s t-test, n = 3). **d** Growth was assessed by cumulative population doublings (cPD) in early passage HCF cells transfected with let-7b-5p and control mimics. **e** SA-β-gal activity and proliferation were analyzed in HCF cells overexpressing let-7b-5p by SA-β-gal and EdU staining. Left: Images of SA-β-gal and EdU stained control (siNeg) and H19KD (siH19) HCF cells are shown. The scale bar represents 200 µm. Right: Quantified data for SA-β-gal activity and EdU staining is represented as a percentage of positive cells (Mean±SEM, *p < 0.05 and ***p < 0.001 by two-tailed unpaired Student’s t-test, n = 3). **f** Senescence-associated secretory phenotype (SASP) in cells transfected with let-7b-5p and control mimics was assessed by RT-qPCR (Mean±SEM, *p < 0.05 and **p < 0.01 by two-tailed unpaired Student’s t-test, n = 3). Source data are provided as a Source Data file.

To further determine how let7b levels contribute to a senescence state, we sought to identify major senescence-associated targets of let7b. Analysis using TargetScan 6.0 indicated specific let7b binding sites in the 3′UTR of EZH2 mRNA **(Figure 5a)**. Interestingly EZH2 has been described as a significant regulator of the senescence program (68, 69). As the enzymatic component of the Polycomb Repressive Complex 2 (PRC2), EZH2 catalyzes the trimethylation of lysine 27 on histone H3 (H3K27me3), resulting in transcriptional repression of genes including p16^INK4A^, p21^Cip1/Waf1^, and other senescence-associated genes. Down-regulation of EZH2 is often associated with cellular senescence, which is dictated by the loss of the EZH2-driven H3K27me3 repressive marks at specific senescence gene loci (69). Consistent with the reported role in senescence, EZH2 mRNA and protein levels were reduced in late passage cells but were maintained by rapamycin treatment **(Figure 5b and c)**. The link between EZH2 and senescence in HCFs is further supported by the growth cessation of early passage HCF cells treated with the EZH2 inhibitor GSK343 **(Figure 5d)**. Similar to the results following targeting of H19, rapamycin was unable to rescue senescence caused by EZH2 targeting, suggesting a causal relationship. Next, using RT-qPCR and western blot analysis, we confirmed that endogenous EZH2 mRNA and protein are indeed suppressed in early passage HCF cells transfected with synthetic let7b mimics **(Figure 5e and f)**. Finally, we confirmed that EZH2 was also reduced following H19 depletion, CTCF depletion, and Nutlin-3 treatment, demonstrating that in each case, EZH2 appears to be a downstream mediator of senescence **(Figure 5g and h, Figure 5-figure supplement 5a-d)**. Because a critical function of EZH2 in the senescence program is suppression of the CDKN2A promoter, we examined the degree of methylation following H19 depletion using Cut and Tag sequencing. We observed decreased binding of an H3K27me3 specific antibody at the CDKN2A consistent with a reduction in methylation of the region and increased expression of p16^INK4A^. A global evaluation of H3K27me3 from this analysis reveals regions of the genome that exhibit generally reduced binding, which suggests the loss of EH2 results in a general decrease in PRC2 activity (**Figure 5-figure supplement 5a-d)**.

**Fig 5:**
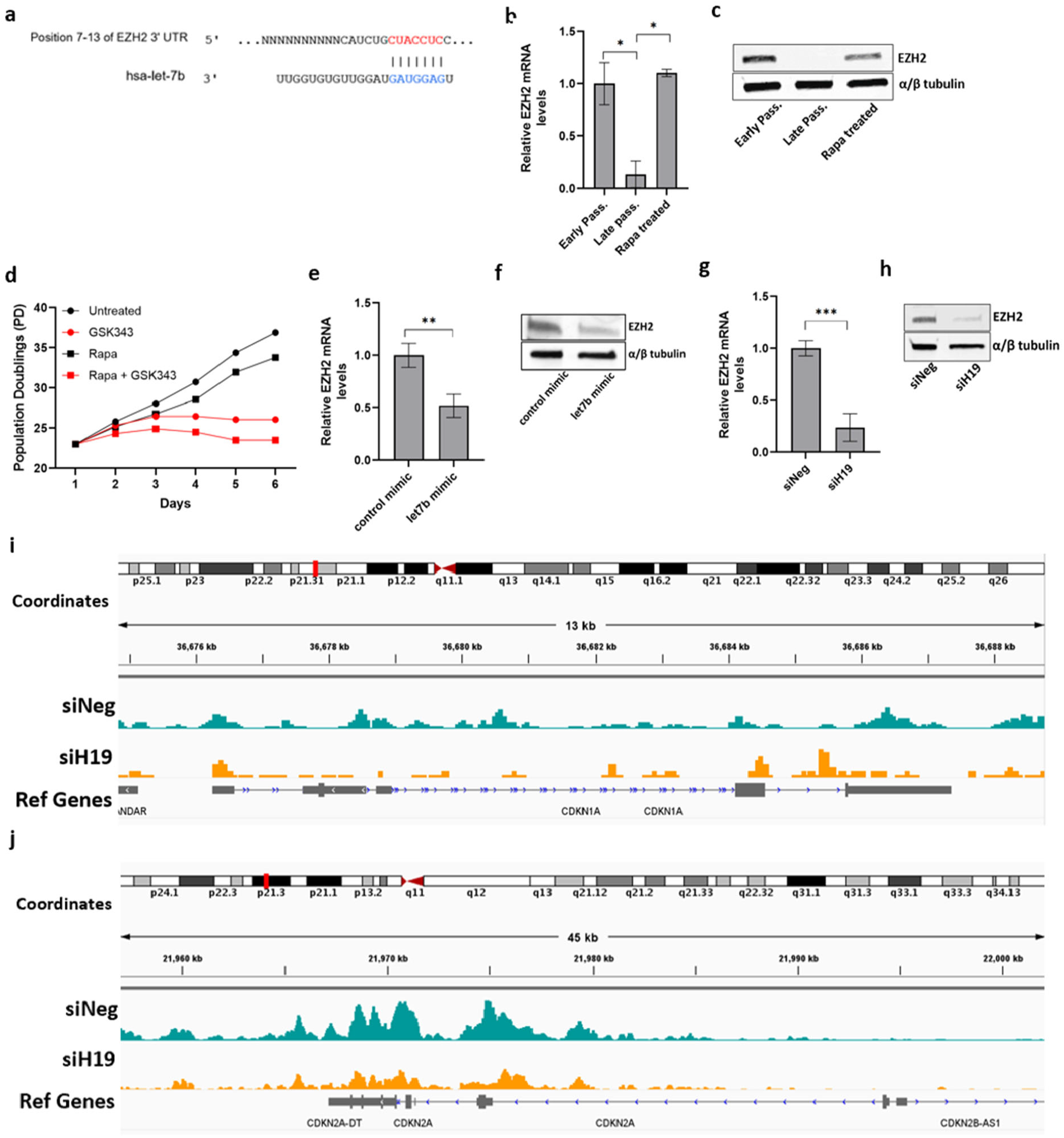
let-7b/EZH2 axis in H19 depletion and senescence. **a** Let-7b binding sites in the 3’UTR of EZH2 mRNA as predicted by TargetScan 6.0. **b** EZH2 mRNA expression was determined by RT-qPCR in the early passage, late passage, and late passage human cardiac fibroblasts (HCF) treated with rapamycin (Mean±SEM, *p < 0.05 by two-tailed unpaired Student’s *t*-test, n = 3). **c** EZH2 protein expression was analyzed by western blot in the early, late, and late passage human cardiac fibroblasts (HCF) treated with rapamycin. α/β tubulin serves as the loading control. **d** Early passage cells grown with or without rapamycin-containing media were treated with an EZH2 inhibitor, GSK343, and growth was assessed by cumulative population doublings (cPD). **e** Early passage cells were transfected with control or let-7b mimic, and EZH2 mRNA expression was determined by RT-qPCR (Mean±SEM, **p < 0.01 by two-tailed unpaired Student’s *t*-test, n = 3). **f** EZH2 protein expression was analyzed in early passage cells transfected with control or let7b overexpression vector, and EZH2 mRNA expression was determined by RT-qPCR. **G** Early passage HCF cells were transfected with siRNA targeting H19 (siH19) and negative control (siNeg), and EZH2 mRNA were determined 7 days after transfection (Mean±SEM, ***p < 0.001 by two-tailed unpaired Student’s *t*-test, n = 3). **H** EZH2 protein levels were analyzed in siH19 and siNeg transfected cells. **I** Representative tracks from H3K27me3 CUT&Tag-sequencing for p21^Cip1/Waf1^ (CDKN1A) gene. **J** Representative tracks from H3K27me3 CUT&Tag-sequencing for p16^INK4A^ (CDKN2A) gene. Tracks show H3K27me3 marks at the p16^INK4A^ gene for early and late passage HCF cells in teal and yellow, respectively. The Refseq gene track is displayed in grey. Source data are provided as a Source Data file.

**Fig 6:**
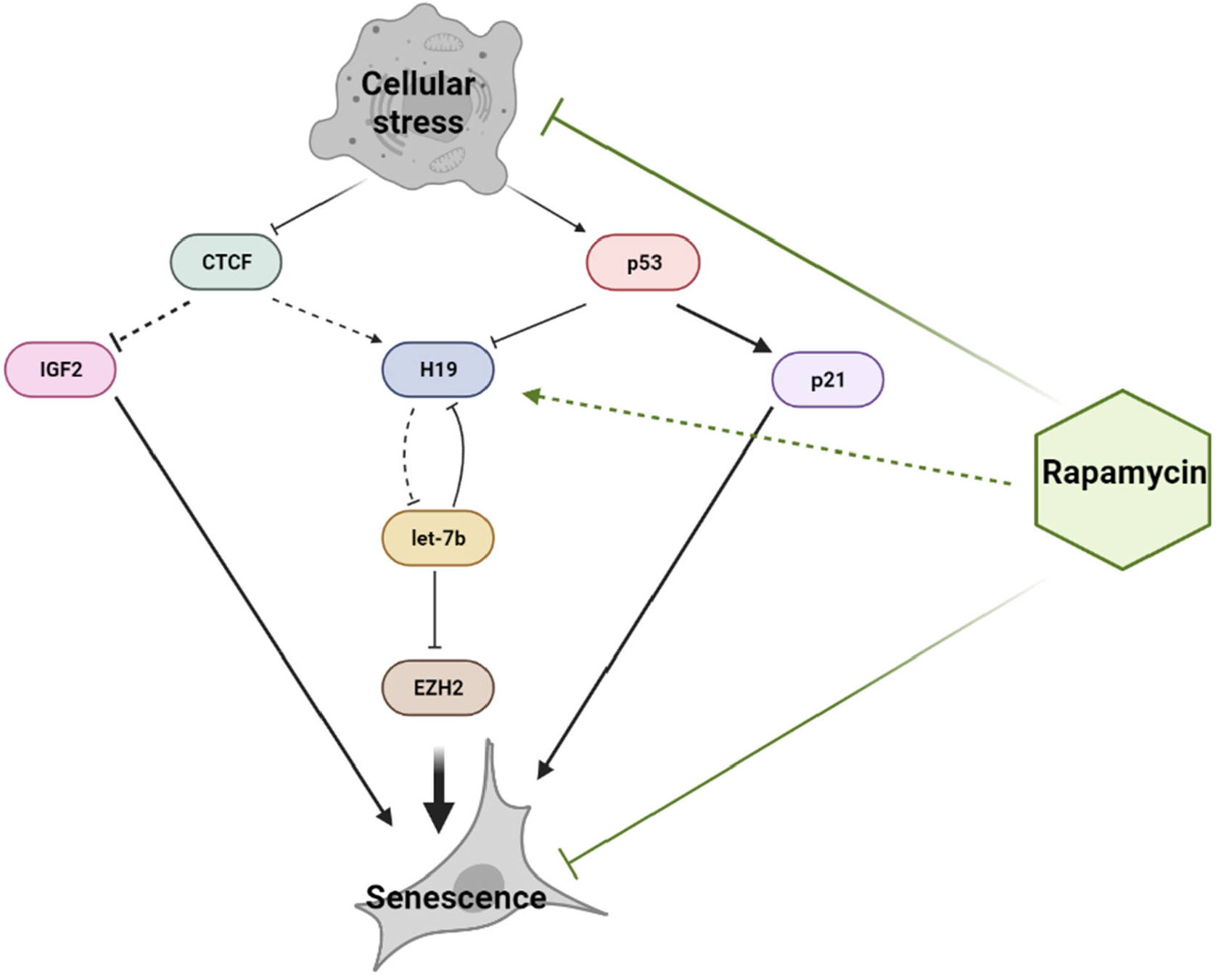
lncRNA H19/let7b/EZH2 axis in senescence. Schematic of the H19/let7b/EZH2 axis in senescing shows that stress-induced loss of CTCF and activation of p53 leads to a reduction in lncRNA H19 levels. Decreased H19 levels result in increased let7b availability and subsequently increased let7b mediated degradation of EZH2. Downregulation of EZH2 expression induces premature senescence. Additionally, increased IGF2 and p21^Cip1/Waf1 expression^ feed into the senescence program. Rapamycin delays the onset of senescence by maintaining lncRNA H19 levels.

## Discussion

In summary, we demonstrate that induction of senescence in somatic cells is mediated by a reduction in lncRNA H19 levels. We propose that the decrease in H19 expression is driven by two convergent pathways: Loss of CTCF expression and the prolonged activation of p53. The loss of H19 further triggers the induction of the senescence program by promoting let7b mediated targeting of EZH2, a critical component in the regulation of senescence-associated genes. More importantly, prolonged treatment with mTOR inhibitor rapamycin maintains lncRNA H19 levels by preventing the loss of CTCF expression and activation of p53, thus preventing the induction of senescence. Taken together, these observations suggest that H19 acts as a critical node in a network that maintains the balance between healthy somatic cell growth and the onset of senescence **(Figure 5)**. Interestingly, such a model can be applied to the growth of stem cells, where H19 is known to have implications for stem cell proliferation and differentiation (70-72). Notably, our data indicate that H19 is required for senescence delaying effects of rapamycin. Thus, H19, which governs two essentially opposing pathways of proliferation and senescence, could serve as a promising target for longevity-enhancing interventions.

CTCF is a critical effector of somatic cell viability and proliferation and is downregulated during senescence (48-50). Notably, some findings identify the role of CTCF in regulating the homologous recombination-directed DNA double-strand break repair (73, 74) and facilitating telomeric replication during cellular stress (75). However, in the case of cardiac fibroblasts subjected to overwhelming damage, CTCF downregulation is not an immediate event but appears to be a gradual progression underlying replicative senescence. An event that immediately follows acute cellular damage is the activation of p53 via the DDR pathway orchestrated by ATM (76). Our data evidently shows that both the downregulation of CTCF and the activation of p53 contribute to the depletion of H19 levels. The mechanisms that lead to decreased H19 expression as part of the senescence program consist of two phases: an acute response driven by p53 activation and a prolonged response dictated by the loss of CTCF. As part of the senescence cell fate decision, these pathways as further accompanied by the activation of p21^Cip1/Waf1^ and upregulation of IGF2, which lie downstream of activation of p53 and CTCF depletion **(Figure 5)**.

While we demonstrate the significance of the H19/let7b/EZH2 axis in the context of senescence in fibroblasts, the vast repertoire of cell and tissue-specific let7b targets indicates that the H19 could potentially regulate other targets of let7b, including HMGA2, Nrf2, E2F2, and c-myc contributing to senescence (77-79). Although we identify let7b as the primary downstream target of lncRNA H19 in human cardiac fibroblasts, most let7 family members can effectively bind to lncRNAH19 due to the conserved canonical and non-canonical lncRNA H19 binding sites shared by the let7 family members (60). Thus, H19 could have additional cellular functions beyond let7b and the proposed axis, creating regulatory networks that govern multiple cell fate decisions.

These regulatory networks involving H19 are likely to impact several cellular processes. For example, evidence suggests that H19 interacts with and regulates the recruitment of methyl-CpG-binding domain protein 1 (MBD1) at target genes, including the IGN network (80). While we show that H19 regulates EZH2 via let-7b, other studies have identified a direct regulation of EZH2 by lncRNA H19 dictated by an interaction between the two (81-83). Further, lncRNA H19 regulates TET1, a DNA demethylase that is responsible for the oxidative conversion of 5-methylcytosine (5mC) to 5-hydroxymethylcytosine (5hmC), which is crucial in the process of base excision repair (84). Together, this creates a picture where lncRNA H19 is at the center of a nexus that regulates multiple aspects of cell growth and maintenance by modulating the network of miRNAs, epigenome modifiers, and senescence-associated genes. Given the emerging picture of senescence as a feature of organismal aging, we anticipate that a greater understanding of H19 regulation of the senescence program will provide insight into age-related diseases and may provide a marker for therapeutic interventions and a potential new target.

## Materials and Methods

### Cell culture

Human Cardiac Fibroblast (HCF) cells were cultivated in Minimal Essential Medium (MEM) supplemented with 10% fetal bovine serum, L-glutamine, MEM vitamins, pyruvate, and MEM nonessential amino acids according to a standard culture protocol for lifespan analysis of human diploid fibroblasts (85). Parallel sets of cultures were maintained in an identical growth medium with the addition of 1 nM Rapa (Enzo Life Sciences). Immortalized Li Fraumeni skin fibroblasts were cultured as described in Medcalf *et al*., 1996 (86). Sequencing genomic DNA obtained from these cells was used to confirm the p53 mutation status (TP53 R175h mutation). HEK-293 T cells for viral production were maintained in DMEM containing 2 mM l-glutamine, 110 mg/mL sodium pyruvate, and 10% FBS (GemCell). For treatment with damaging agents, cells were exposed to Neocarzinostatin (NCS) (0.2μM) (Bristol Lab) for 2hrs and H2O2 (200μM) (H325-500, Fisher Chemicals) for 1hr. RNA and protein extraction was performed 24hrs post-treatment. For p53 activation, cells were exposed to MDM2 inhibitor, Nutlin-3 (444143, Sigma-Aldrich) for 72hrs. Pharmacological inhibition with EZH2 inhibitor GSK343 (A3449, APEXBIO) (5μM) was maintained across the lifespan as indicated.

### Viral infection

To produce the shp53 knockdown HCF cells, we used a pLKO.1 lentiviral vector (MISSION^®^ TRC shRNA TRCN0000003756) containing either a p53 short hairpin RNA or a scrambled shRNA sequence (Addene Cat# 1864). 10g of plasmid DNA was transfected into 293T lentiviral packaging cells using LT1 transfection reagent (Mirus Bio LLC), and viral supernatants were collected. Early passage HCF cells were infected with p53 shRNA or scrambled shRNA viral supernatant and 8 µg/mL polybrene (American Bioanalytical) for 24 hours, followed by a 72 hour selection with 1 µg/mL puromycin (Mediatech).

### RNA interference assay

According to the manufacturer’s protocol, siRNA against H19, CTCF, and non-targeting control (100 nM) were transfected into cells using the Lipofectamine RNAiMAX (Invitrogen). The siRNA targeting sequences are listed in **Table 1**.

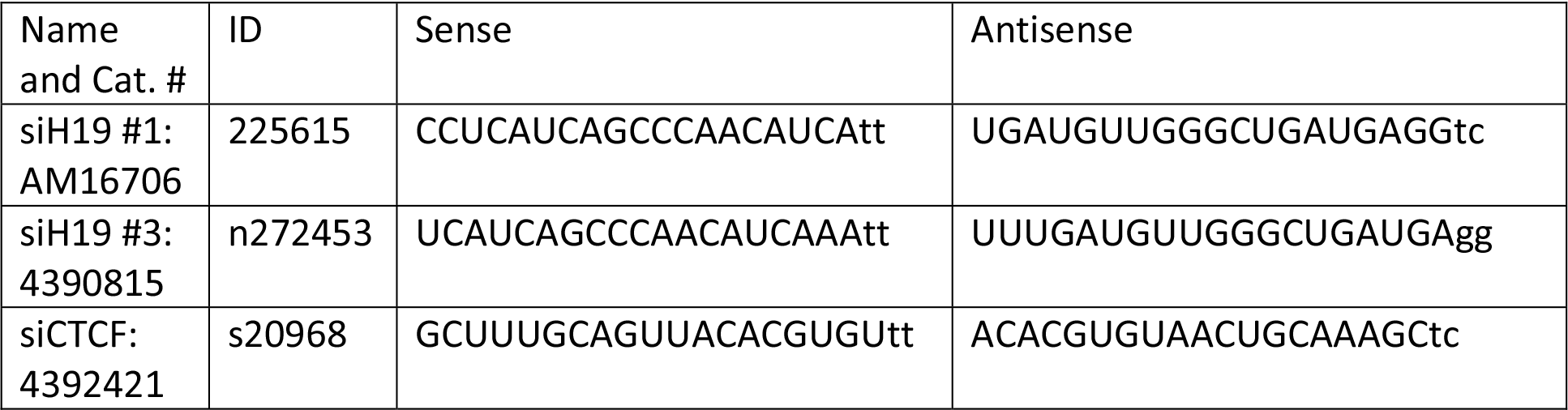

### Let-7b mimics

According to the manufacturer’s protocol, mirVana™ miRNA Mimic, Negative Control #1 (4464058, ThermoFisher) (50 nM), and mirVana^®^ hsa-let7b-5p miRNA mimic (MC24132, ThermoFisher) (50 nM) were transfected into cells using the Lipofectamine RNAiMAX (Invitrogen). Cells were analyzed 96 hrs post-transfection.

### Quantitative real-time PCR (RT-qPCR) and NanoString analysis

RNA was isolated using TRIzol solution (Thermo Fisher Scientific) and quantitative real-time PCR was performed using TaqMan Assays (Thermo Fisher Scientific) (GAPDH: Hs02786624_g1; H19: Hs0026142_g1 and Hs00399294_g1; IGF2: Hs01005964_g1; CTCF: Hs00902016_m1; miR-let7b: 478576_mir; EZH2:vHs00544830_m1; p21: Hs00355782_m1; CXCL8: Hs00174103_m1; MMP3: Hs00968305_m1; CCL5: Hs 99999048_m1; IL-6: Hs 00174131_m1; FGF2: Hs0026645_m1; IL-1A: Hs 00174092_m1) using an ABI™7500 Real-Time system. All samples were analyzed in triplicates. Fold changes were calculated from raw data using the ΔΔCT method. GAPDH was used as an internal control. NanoString analysis was performed according to the manufacturer’s protocol (87)

### Western blot

Cells were lysed in RIPA lysis buffer containing a protease inhibitor cocktail (Sigma-Aldrich), and protein quantification was performed using a BCA assay (Thermo Fisher Scientific). Lysates were run on an SDS-4-15% polyacrylamide gel (Bio-Rad) and transferred onto a Nitrocellulose membrane (Amersham). Immunoblots were visualized with the Odyssey blot imager (LI-COR). Antibodies used were CTCF (D31H2) XP(R) Rabbit monoclonal antibody (Cat. 3418S, Cell Signaling Tech; 1:1000 dilution), EZH2 (D2C9) XP(R) Rabbit monoclonal antibody (Cat. 5246S, Cell Signaling Tech; 1:1000 dilution), p21Cip1/Waf1 Waf1/Cip1 (12D1) Rabbit monoclonal antibody (Cat.2947S, Cell Signaling Tech; 1:1000 dilution), p16^INK4A^ (JC8) mouse monoclonal antibody (Cat.sc-56330, Santa Cruz; 1:1000 dilution), p53 monoclonal antibody (Cat. 39554, ActiveMotif; 1:1000 dilution), α/β tubulin Rabbit antibody (Cat. 2148S, Cell Signaling Tech; 1:1000 dilution), GAPDH (D16H11) XP(R) Rabbit monoclonal antibody (Cat. 5174S, Cell Signaling Tech; 1:1000 dilution)

### Senescence-associated (SA) β-galactosidase staining

Cells were washed twice with PBS and then fixed with 3% formaldehyde for 3 min at room temperature. Cells were then washed twice with PBS and stained with X-gal (Tenova X1220) (1 mg/mL of X-gal in 40 mM citric acid/Na2HPO4 [pH 6], 5 mM potassium ferrocyanide, 5 mM potassium ferricyanide, 150 mM NaCl, 2 mM MgCl2) for 18 h at 37 °C. Cells were washed with PBS twice and imaged using EVOS FL Auto microscope (Thermo Fisher). The percentage of positively stained cells was determined by counting seven random fields. Images of representative fields were captured under 40X magnification.

### EdU labeling

Cells were washed twice with PBS and then fixed with 3% formaldehyde for 10 min at room temperature. Cells were washed twice with PBS and permeabilized with 0.1% Triton/TBS for 10 min. Cells were incubated with EdU stain (100mM Tris (pH8.5), 1mM CuSO_4_, 1.25 μM Azide Fluor 488, and 50mM ascorbic acid) at room temperature for 30 mins. Cells were washed with PBS twice and imaged using EVOS FL Auto microscope (Thermo Fisher). The percentage of positively stained cells was determined by counting seven random fields. Images of representative a were captured under 20X magnification.

### Cleavage Under Targets and Tagmentation sequencing (CUT&Tag-sequencing)

CUT&Tag was performed using CUT&Tag-IT™ Assay Kit (53160, ActiveMotif) according to the manufacturer’s instructions using 5 × 105 cells. The raw reads were processed and mapped using BasePair. For each obtained sample, adaptors were trimmed. The trimmed reads were aligned to the GRCh38 human reference genome using Bowtie2. Duplicate fragment reads and reads mapping to the mitochondrial genome were excluded from all downstream analyses. Gene body coverage was computed using Galaxy tools (Afgan et al., 2018) (https://usegalaxy.org/).

### Methylation Analysis

Genomic DNA was isolated using a DNA purification kit from Zymo Scientific according to the manufacturer’s directions. DNA was subjected to bisulfite modification using the Epimark bisulfite conversion kit from New England Biolabs. CTCF binding site 5 was amplified using primers described by Takai et al. (56). PCR products were cloned using the Topo TA cloning kit (Invitrogen). Individual colonies were propagated, and plasmid DNA was sequenced by Sanger Sequencing (GeneWiz).

### Statistical Analysis

Data are represented as mean ± SEM or mean ± SD as described in the figure legends. The differences between groups were determined by two-tailed Student’s *t-test, P* values are indicated by non-significant (p > 0.05), *(p < 0.05), **(p < 0.01), or ***(p < 0.001), ****(p<0.0001). Data without an explicit indication of statistical significance should be considered to have a p-value greater than 0.05. All experiments were performed in triplicate, as specified in the figure legends.

## Data availability

The source data for Figs. 1a-h, 2a-e, 3a-f, 4a-f, 5a-j and Supplementary Figs. 1a-g, 2a-e, 3a-c, 4a-b, and 5a-f are provided as a Source Data file.

## Contributions

M.P. and C.S. conceived, designed, conducted the study, analyzed data, and prepared the manuscript. E.N. recommended experiments and edited the manuscript. J.D. and O.E. performed experiments, which are included in the paper.

## Ethics declarations

CS has an equity share in Boinica Therapeutics LLC and is a named inventor on patent US11179374B2.

## Supplementary information

**Supplementary Fig 1:**
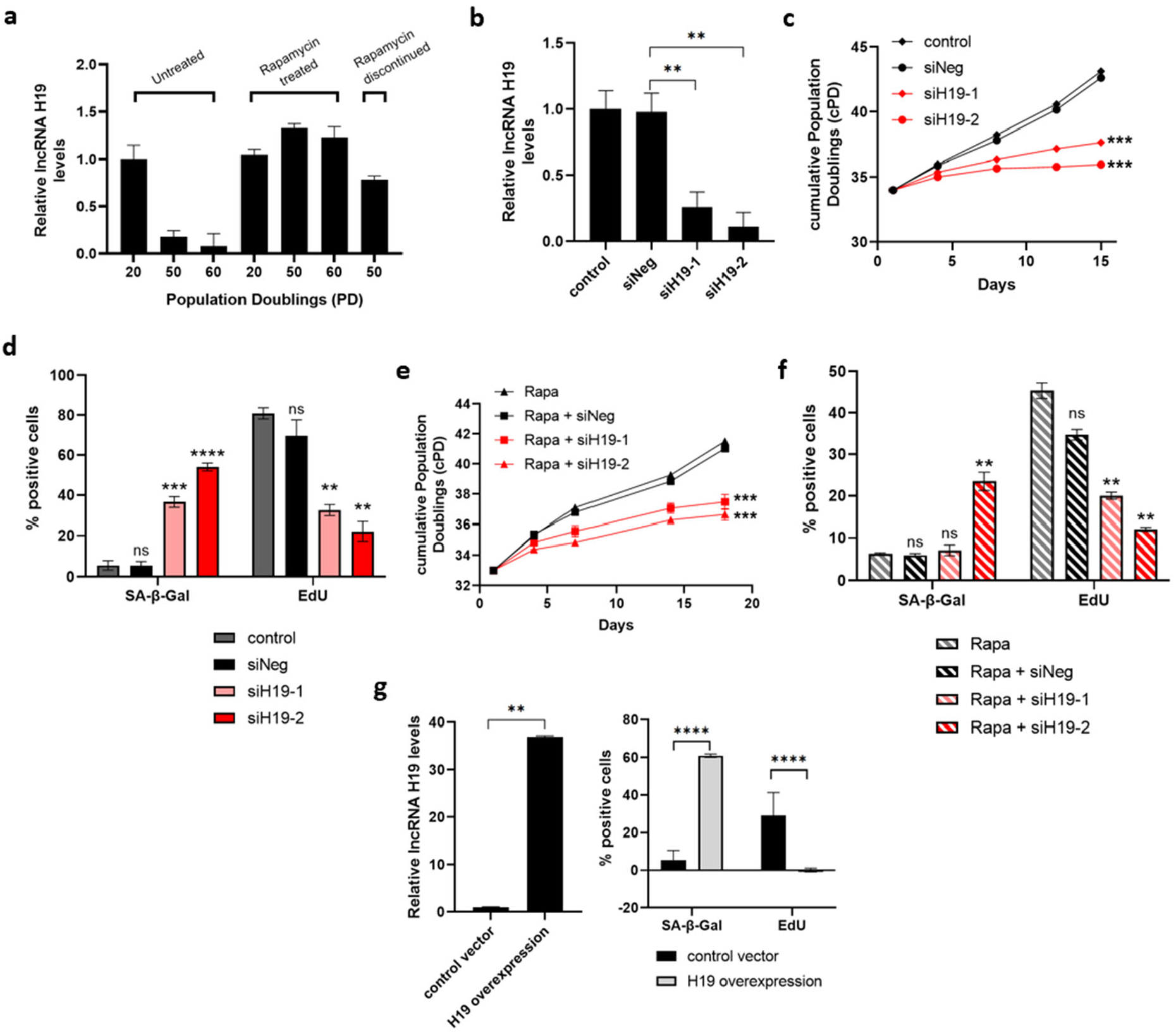
H19 expression is reduced during senescence but maintained with rapamycin treatment. **a** RT-qPCR determined lncRNA H19 steady-state levels in the human cardiac fibroblasts (HCF) at PD 20, 50, and 60 cultured with or without rapamycin-containing media. Rapamycin treatment was terminated at PD 48 to assess H19 levels in rapamycin discontinued PD 50 cells. **b** Early passage HCF cells were transfected with two different siRNAs targeting H19 (siH19-1 and siH19-2) and negative control (siNeg), and levels of lncRNA H19 were determined 4 days after transfection. Control cells represent non-transfected early passage HCF cells (Mean±SEM, **p < 0.01 by two-tailed unpaired Student’s *t*-test, n = 3). **c** Cell growth in H19 targeted cells was assessed by cumulative population doublings (cPD) (Mean±SEM, ***p < 0.001 by two-tailed unpaired Student’s *t*-test, n = 3). **d** SA-β-gal activity and proliferation were analyzed in H19 knockdown cells by SA-β-gal and EdU staining, respectively. Quantification is represented as a percentage of positive cells (Mean±SEM, p > 0.05, **p < 0.01, ***p < 0.001, and ****p < 0.0001 by two-tailed unpaired Student’s *t*-test, n = 3). **e** Cell growth in rapamycin-treated H19 targeted cells was assessed by cumulative population doublings (cPD) (Mean±SEM, ***p < 0.001 by two-tailed unpaired Student’s *t*-test, n = 3). **f** SA-β-gal activity and proliferation were analyzed in rapamycin-treated H19 knockdown cells by SA-β-gal and EdU staining, respectively. Quantification is represented as a percentage of positive cells (Mean±SEM, p > 0.05 and **p < 0.01 by two-tailed unpaired Student’s *t*-test, n = 3). **g** Early passage HCF cells were infected with H19 overexpressing lentiviruses, and levels of lncRNA H19 were determined after infection. SA-β-gal activity and proliferation were analyzed in H19 overexpressing HCF cells by SA-β-gal and EdU staining. Left: lncRNA H19 levels as measured with RT-qPCR. Right: Quantification is represented as a percentage of positive (Mean±SEM, ****p < 0.0001 by two-tailed unpaired Student’s *t*-test, n = 3). Source data are provided as a Source Data file.

**Supplementary Fig 2:**
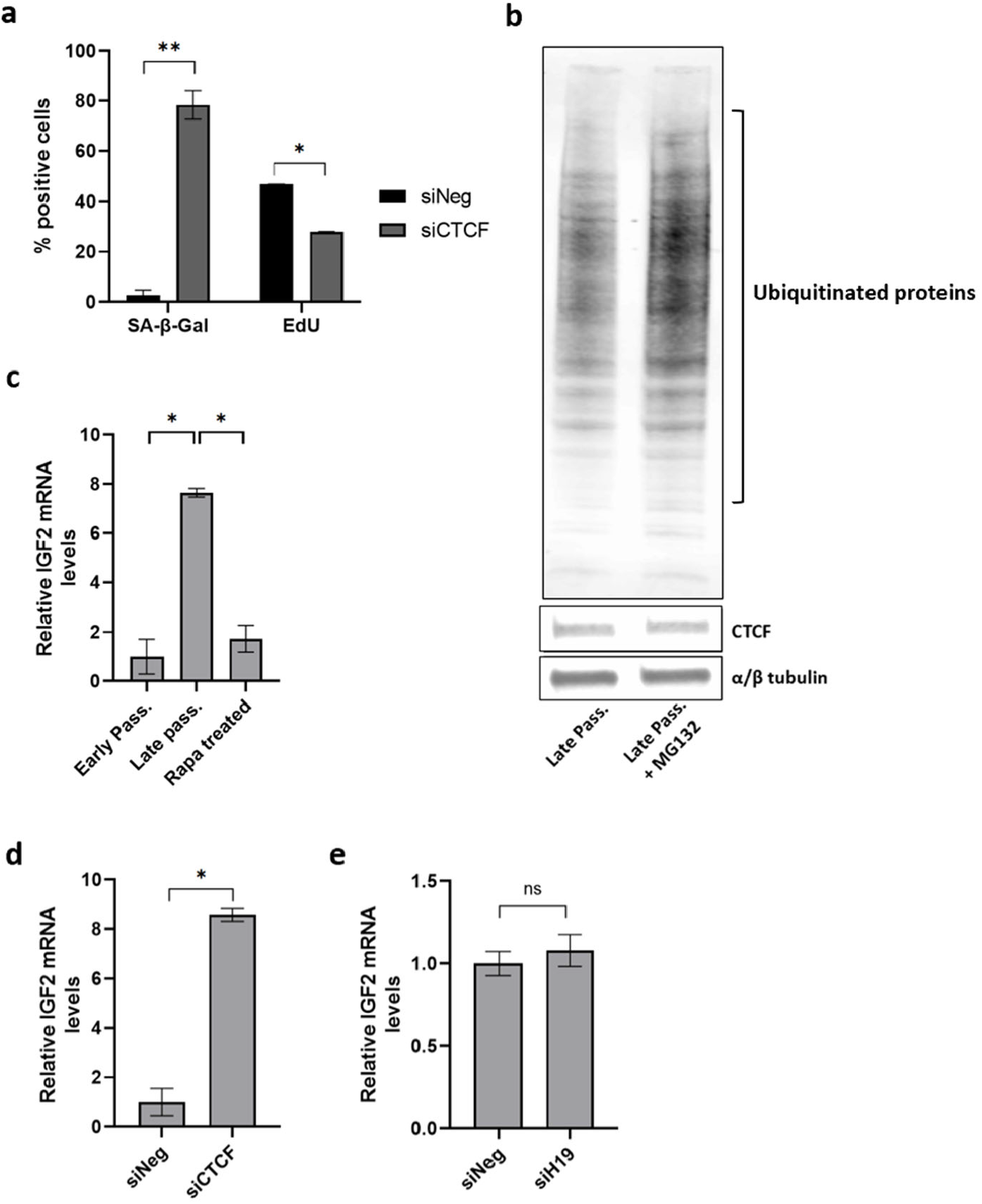
Loss of CTCF leads to a reduction in H19 expression. **a** SA-β-gal activity and proliferation were analyzed in siCTCF cells by SA-β-gal and EdU staining. Quantified data is represented as a percentage of positive cells (Mean±SEM, *p < 0.05 and **p < 0.01 by two-tailed unpaired Student’s *t*-test, n = 3). **b** CTCF protein expression was analyzed by western blot in the late passage and late passage HCF cells treated with a proteasome inhibitor (MG132). Total ubiquitinated proteins were analyzed to confirm the effectiveness of MG132. α/β tubulin serves as the loading control. **c** IGF mRNA levels were determined by RT-qPCR in the early passage, late passage, and late passage HCF cells treated with rapamycin (Mean±SEM, *p < 0.05 by two-tailed unpaired Student’s *t*-test, n = 3). **d** Early passage HCF cells were transfected with siRNA targeting CTCF (siCTCF) and negative control (siNeg), and IGF mRNA levels were determined by RT-qPCR (Mean±SEM, *p < 0.05 by two-tailed unpaired Student’s *t*-test, n = 3). **e** IGF mRNA levels were determined by RT-qPCR in the early passage HCF cells transfected with siRNA targeting H19 (siH19) and negative control (siNeg) (Mean±SEM, p > 0.05 by two-tailed unpaired Student’s *t*-test, n = 3). Source data are provided as a Source Data file.

**Supplementary Fig 3:**
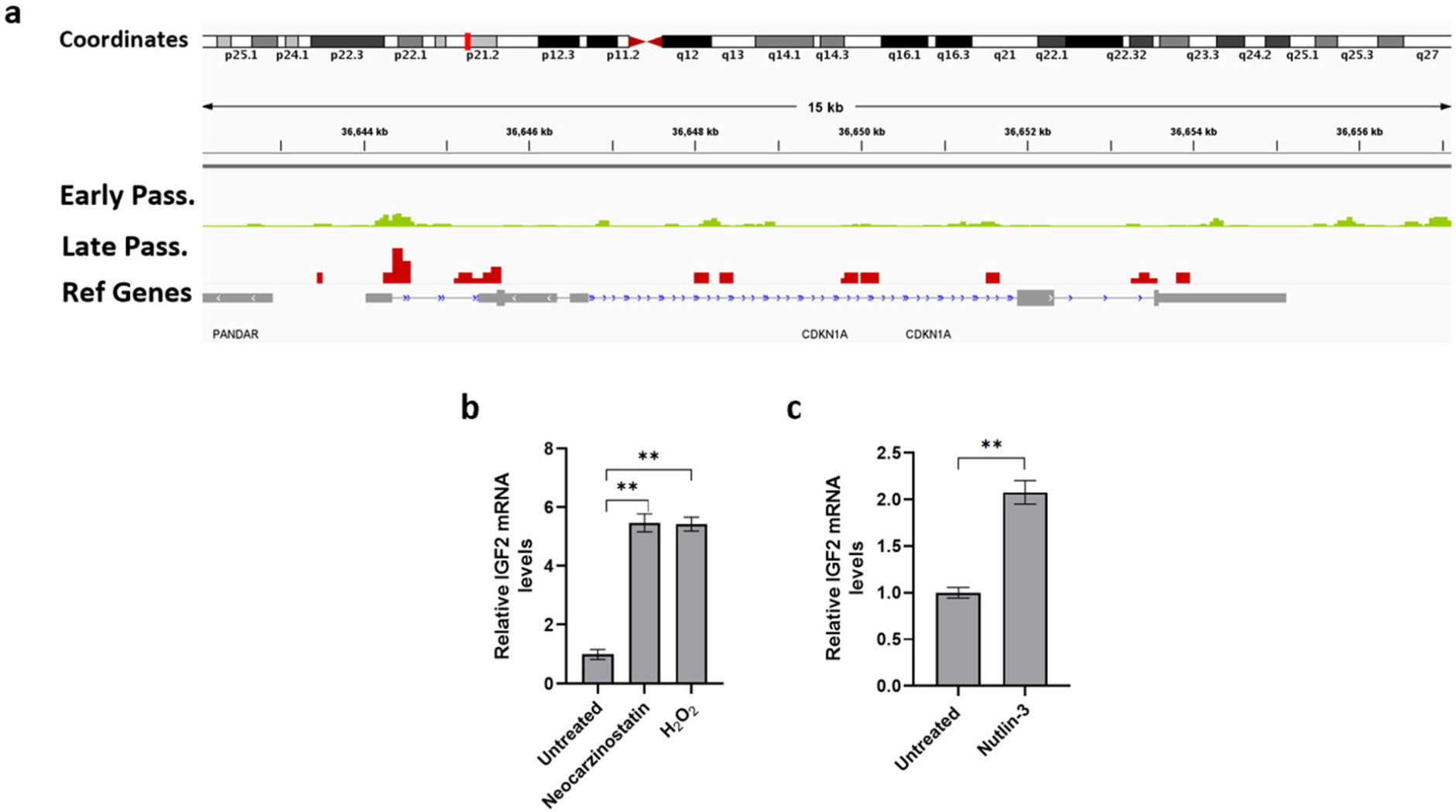
p53 activation results in decreased H19 expression. **a** Representative tracks from p53 CUT&Tag-sequencing for p21^Cip1/Waf1^ (CDKN1A). Tracks show p53 binding at the p21^Cip1/Waf1^ gene for early and late passage HCF cells in green and red, respectively. The Refseq gene track is displayed in grey. **b** Early passage HCF cells were treated with genotoxin neocarzinostatin (0.2μM) and hydrogen peroxide (200μM), and IGF2 mRNA levels were determined 24 hrs after treatment (Mean±SEM, **p < 0.01 by two-tailed unpaired Student’s *t*-test, n = 3). **c** Early passage HCF cells were treated with MDM2 inhibitor Nutlin-3, and IGF2 mRNA levels were determined 24 hrs after treatment (Mean±SEM, **p < 0.01 by two-tailed unpaired Student’s *t*-test, n = 3). Source data are provided as a Source Data file.

**Supplementary Fig 4:**
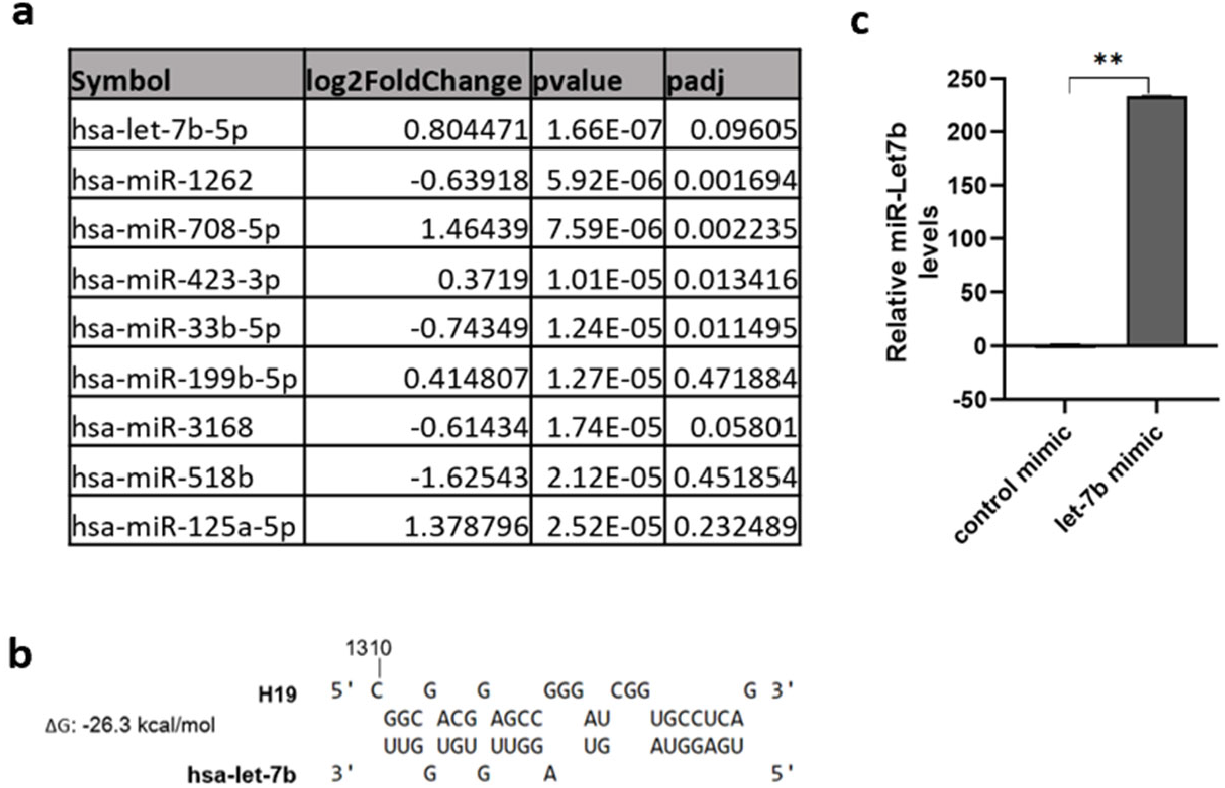
let-7b/EZH2 axis in H19 depletion and senescence. **a** List of top 10 upregulated miRNAs from nCounter^®^ miRNA Expression Panels. **b** Partial sequences of H19 and let-7b as predicted by RNAhybrid - BiBiServ2. **c** Early passage HCF cells were transfected with control and miR-let7b-5p mimic. miR-let7b levels were determined 48hrs after transfection (Mean±SEM, **p < 0.01 by two-tailed unpaired Student’s *t*-test, n = 3). Source data are provided as a Source Data file.

**Supplementary Fig 5:**
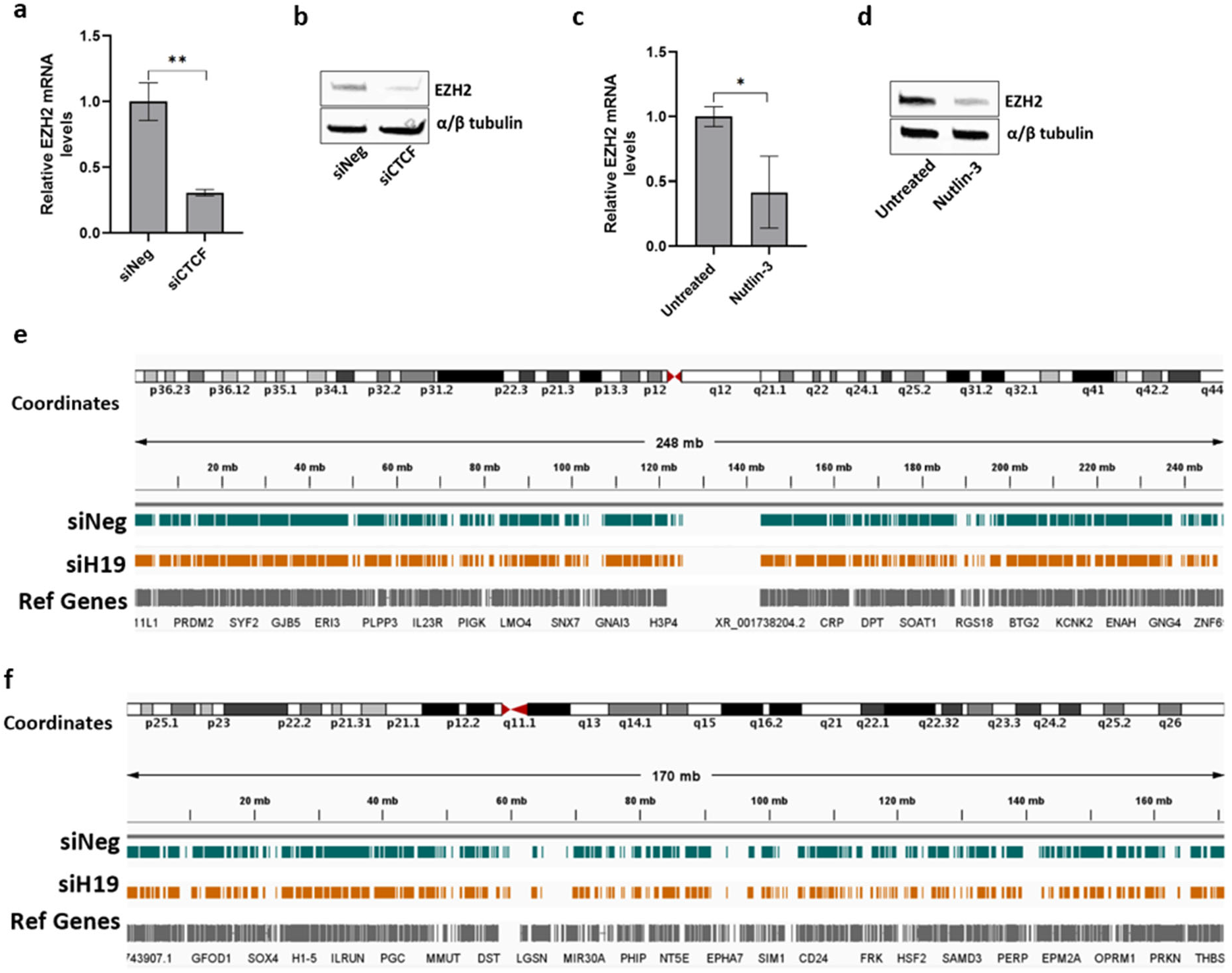
let-7b/EZH2 axis in H19 depletion and senescence. **a** Total RNA from early passage HCF cells transfected with siRNA targeting CTCF (siCTCF) and negative control (siNeg) was subjected to RT-qPCR, and EZH2 mRNA levels were determined 7 days after transfection (Mean±SEM, **p < 0.01 by two-tailed unpaired Student’s *t*-test, n = 3). **b** EZH2 protein expression was analyzed by western blot in cells transfected with siRNA targeting CTCF (siCTCF) and negative control (siNeg). α/β tubulin serves as the loading control. **c** HCF cells were treated with MDM2 inhibitor Nutlin-3, and EZH2 mRNA levels were determined 72 hrs after treatment (Mean±SEM, *p < 0.05 by two-tailed unpaired Student’s *t*-test, n = 3). **d** EZH2 protein expression was analyzed by western blot in cells treated with Nutlin-3 (72hr post-treatment). α/β tubulin serves as the loading control. **e** Representative tracks from H3K27me3 CUT&Tag-sequencing for chromosome 1. **f** Representative tracks from H3K27me3 CUT&Tag-sequencing for chromosome 6. Tracks for control (siNeg) and H19 depleted (siH19) HCF cells are displayed in teal and yellow, respectively. The Refseq gene track is displayed in grey. Source data are provided as a Source Data file.

