## Supplemental material for "lncRNA H19/Let7b/EZH2 axis regulates somatic cell senescence"

Drexel University College of Medicine

245 North 15th Street

Philadelphia, PA 19102

### Supplementary information

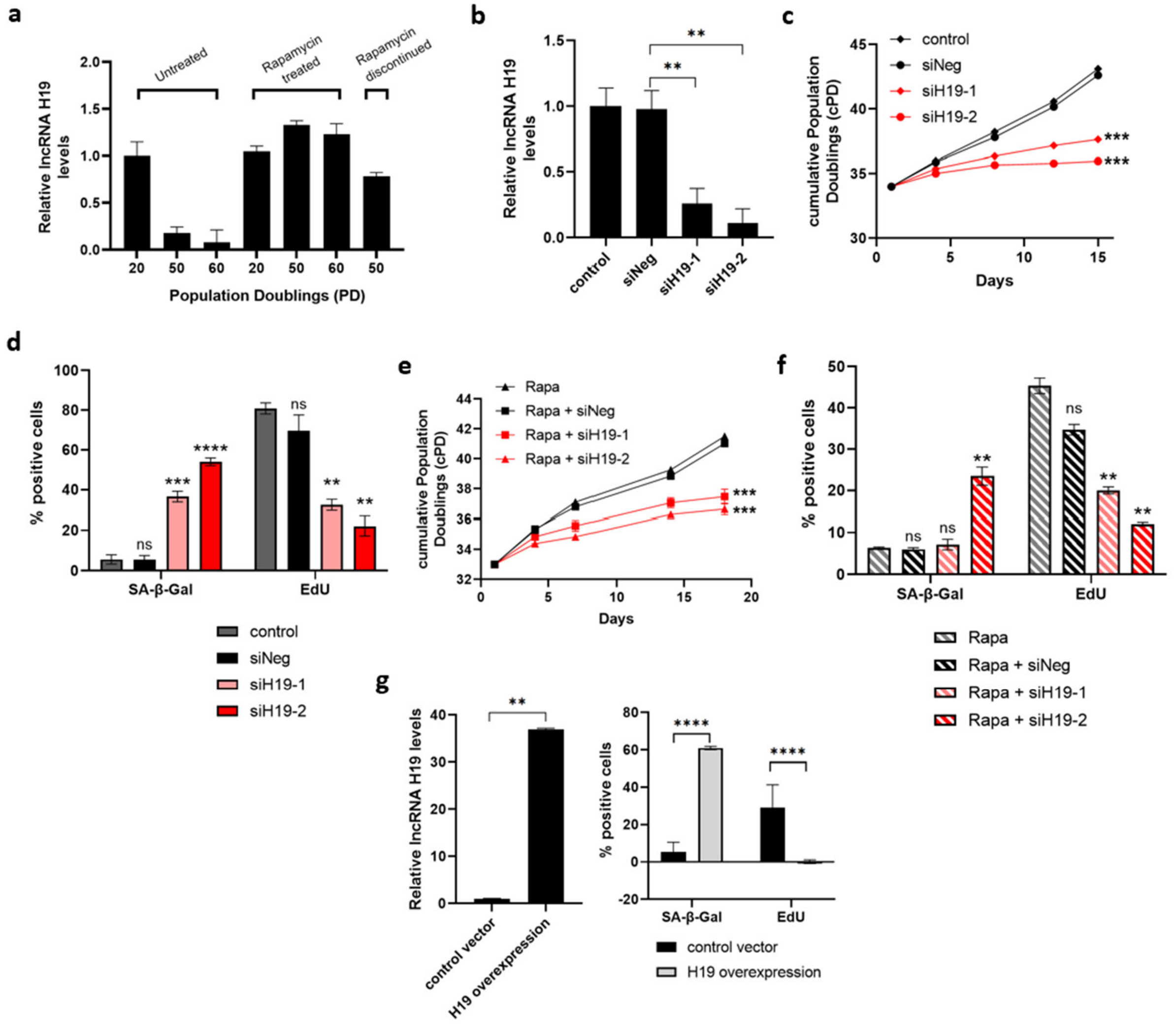


**Supplementary Fig 1: H19 expression is reduced during senescence but maintained with rapamycin treatment**

**a** RT-qPCR determined lncRNA H19 steady-state levels in the human cardiac fibroblasts (HCF) at PD 20, 50, and 60 cultured with or without rapamycin-containing media. Rapamycin treatment was terminated at PD 48 to assess H19 levels in rapamycin discontinued PD 50 cells. **b** Early passage HCF cells were transfected with two different siRNAs targeting H19 (siH19-1 and siH19-2) and negative control (siNeg), and levels of lncRNA H19 were determined 4 days after transfection. Control cells represent non-transfected early passage HCF cells (Mean±SEM, **p < 0.01 by two-tailed unpaired Student's *t*-test, n = 3). **c** Cell growth in H19 targeted cells was assessed by cumulative population doublings (cPD) (Mean±SEM, ***p < 0.001 by two-tailed unpaired Student's *t*-test, n = 3). **d** SA-β-gal activity and proliferation were analyzed in H19 knockdown cells by SA-β-gal and EdU staining, respectively. Quantification is represented as a percentage of positive cells (Mean±SEM, p > 0.05, **p < 0.01, ***p < 0.001, and ****p < 0.0001 by two-tailed unpaired Student's *t*-test, n = 3). **e** Cell growth in rapamycin-treated H19 targeted cells was assessed by cumulative population doublings (cPD) (Mean±SEM, ***p < 0.001 by two-tailed unpaired Student's *t*-test, n = 3). **f** SA-β-gal activity and proliferation were analyzed in rapamycin-treated H19 knockdown cells by SA-β-gal and EdU staining, respectively. Quantification is represented as a percentage of positive cells (Mean±SEM, p > 0.05 and **p < 0.01 by two-tailed unpaired Student's *t*-test, n = 3). **g** Early passage HCF cells were infected with H19 overexpressing lentiviruses, and levels of lncRNA H19 were determined after infection. SA-β-gal activity and proliferation were analyzed in H19 overexpressing HCF cells by SA-β-gal and EdU staining. Left: lncRNA H19 levels as measured with RT-qPCR. Right: Quantification is represented as a percentage of positive (Mean±SEM, ****p < 0.0001 by two-tailed unpaired Student's *t*-test, n = 3). Source data are provided as a Source Data file.

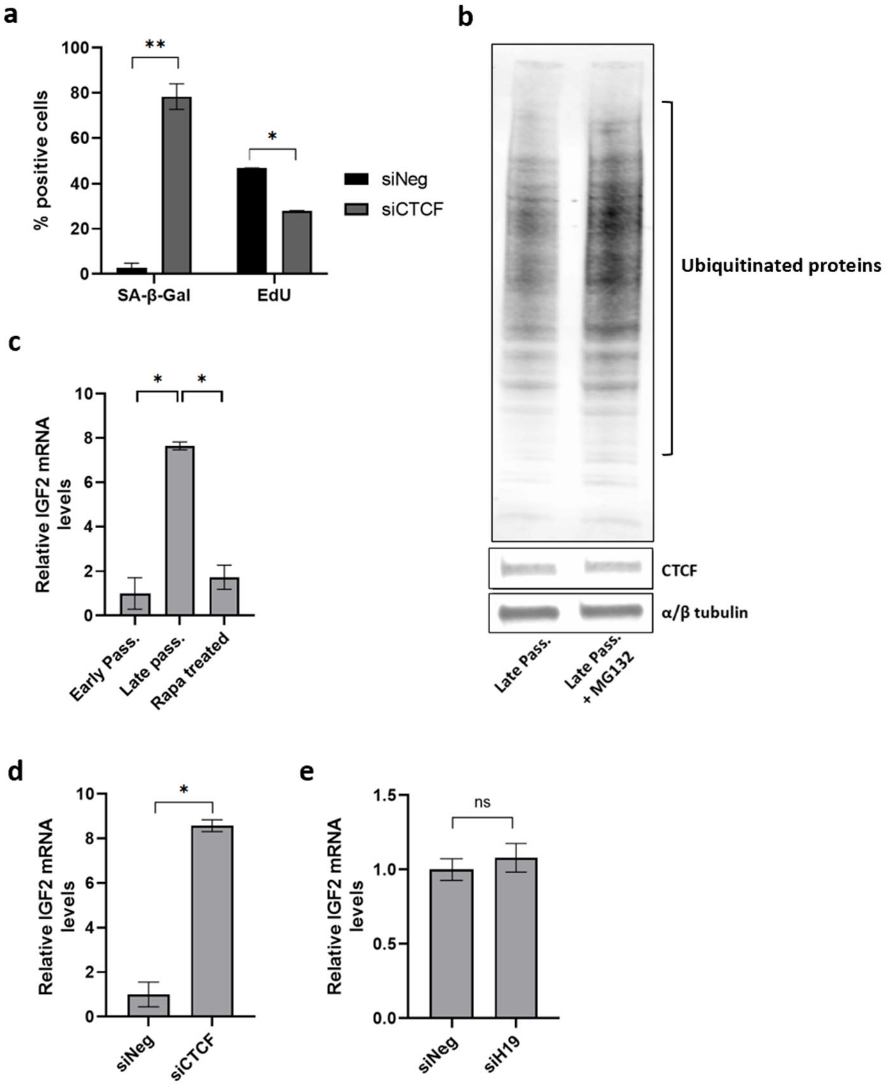


**Supplementary Fig 2: Loss of CTCF leads to a reduction in H19 expression**

**a** SA-β-gal activity and proliferation were analyzed in siCTCF cells by SA-β-gal and EdU staining. Quantified data is represented as a percentage of positive cells (Mean±SEM, *$p < 0.05$ and **$p < 0.01$ by two-tailed unpaired Student's *t*-test, n = 3). **b** CTCF protein expression was analyzed by western blot in the late passage and late passage HCF cells treated with a proteasome inhibitor (MG132). Total ubiquitinated proteins were analyzed to confirm the effectiveness of MG132. α/β tubulin serves as the loading control. **c** IGF mRNA levels were determined by RT-qPCR in the early passage, late passage, and late passage HCF cells treated with rapamycin (Mean±SEM, *$p < 0.05$ by two-tailed unpaired Student's *t*-test, n = 3). **d** Early passage HCF cells were transfected with siRNA targeting CTCF (siCTCF) and negative control (siNeg), and IGF mRNA levels were determined by RT-qPCR (Mean±SEM, *$p < 0.05$ by two-tailed unpaired Student's *t*-test, n = 3). **e** IGF mRNA levels were determined by RT-qPCR in the early passage HCF cells transfected with siRNA targeting H19 (siH19) and negative control (siNeg) (Mean±SEM, $p > 0.05$ by two-tailed unpaired Student's *t*-test, n = 3). Source data are provided as a Source Data file.

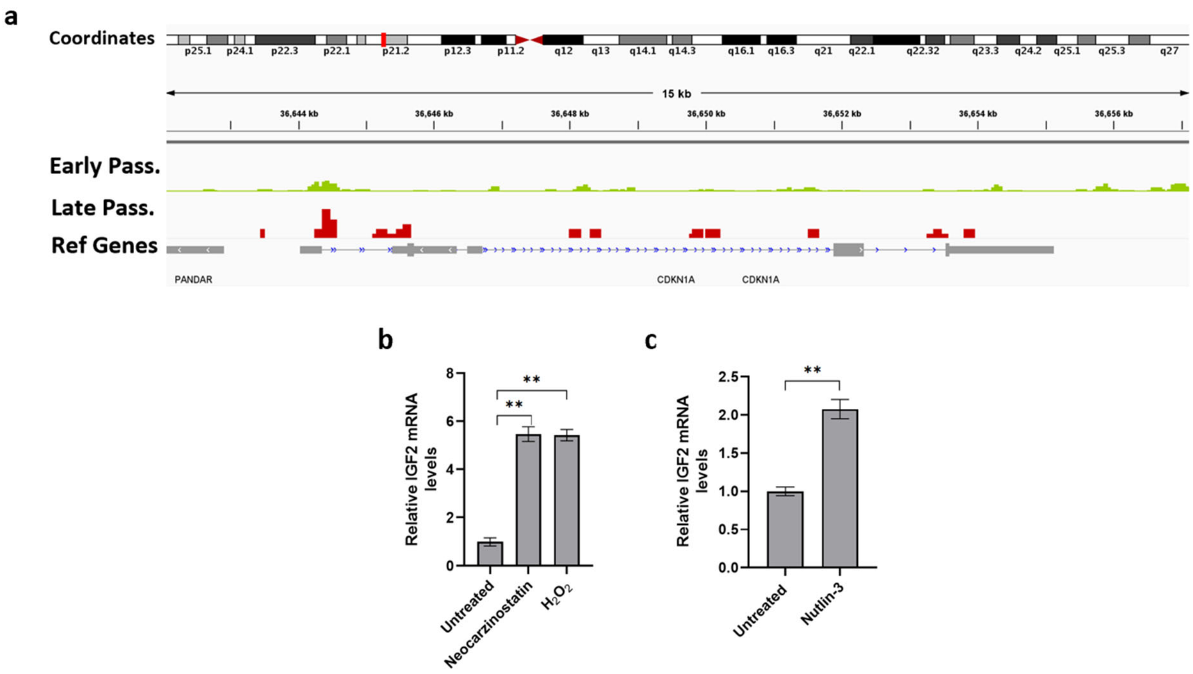


**Supplementary Fig 3: p53 activation results in decreased H19 expression**

**a** Representative tracks from p53 CUT&Tag-sequencing for p21$^{Cip1/Waf1}$ (CDKN1A). Tracks show p53 binding at the p21$^{Cip1/Waf1}$ gene for early and late passage HCF cells in green and red, respectively. The Refseq gene track is displayed in grey. **b** Early passage HCF cells were treated with genotoxin neocarzinostatin (0.2µM) and hydrogen peroxide (200µM), and IGF2 mRNA levels were determined 24 hrs after treatment (Mean±SEM, $^{**}p < 0.01$ by two-tailed unpaired Student's *t*-test, n = 3). **c** Early passage HCF cells were treated with MDM2 inhibitor Nutlin-3, and IGF2 mRNA levels were determined 24 hrs after treatment (Mean±SEM, $^{**}p < 0.01$ by two-tailed unpaired Student's *t*-test, n = 3). Source data are provided as a Source Data file.

**a**

| Symbol | log2FoldChange | pvalue | padj |
|---|---|---|---|
| hsa-let-7b-5p | 0.804471 | 1.66E-07 | 0.09605 |
| hsa-miR-1262 | -0.63918 | 5.92E-06 | 0.001694 |
| hsa-miR-708-5p | 1.46439 | 7.59E-06 | 0.002235 |
| hsa-miR-423-3p | 0.3719 | 1.01E-05 | 0.013416 |
| hsa-miR-33b-5p | -0.74349 | 1.24E-05 | 0.011495 |
| hsa-miR-199b-5p | 0.414807 | 1.27E-05 | 0.471884 |
| hsa-miR-3168 | -0.61434 | 1.74E-05 | 0.05801 |
| hsa-miR-518b | -1.62543 | 2.12E-05 | 0.451854 |
| hsa-miR-125a-5p | 1.378796 | 2.52E-05 | 0.232489 |

**b**

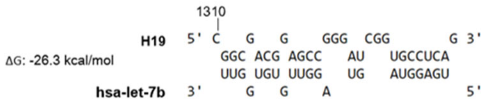


**c**

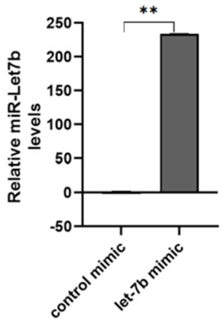


**Supplementary Fig 4: let-7b/EZH2 axis in H19 depletion and senescence.**

**a** List of top 10 upregulated miRNAs from nCounter® miRNA Expression Panels. **b** Partial sequences of H19 and let-7b as predicted by RNAhybrid - BiBiServ2. **c** Early passage HCF cells were transfected with control and miR-let7b-5p mimic. miR-let7b levels were determined 48hrs after transfection (Mean±SEM, **p < 0.01 by two-tailed unpaired Student's *t*-test, n = 3). Source data are provided as a Source Data file.

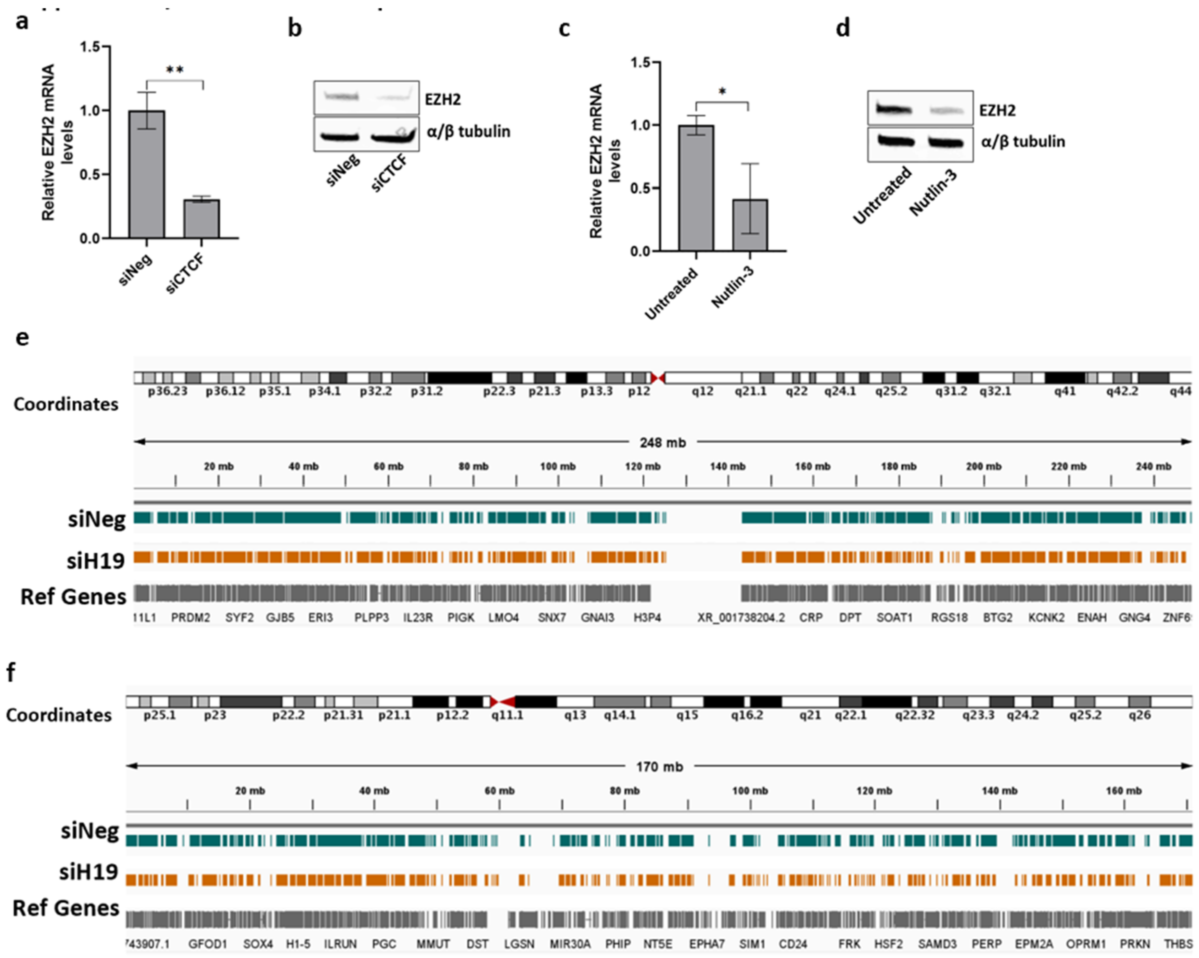


**Supplementary Fig 5: let-7b/EZH2 axis in H19 depletion and senescence.**

**a** Total RNA from early passage HCF cells transfected with siRNA targeting CTCF (siCTCF) and negative control (siNeg) was subjected to RT-qPCR, and EZH2 mRNA levels were determined 7 days after transfection (Mean±SEM, $^{**}p < 0.01$ by two-tailed unpaired Student's *t*-test, n = 3). **b** EZH2 protein expression was analyzed by western blot in cells transfected with siRNA targeting CTCF (siCTCF) and negative control (siNeg). α/β tubulin serves as the loading control. **c** HCF cells were treated with MDM2 inhibitor Nutlin-3, and EZH2 mRNA levels were determined 72 hrs after treatment (Mean±SEM, $^{*}p < 0.05$ by two-tailed unpaired Student's *t*-test, n = 3). **d** EZH2 protein expression was analyzed by western blot in cells treated with Nutlin-3 (72hr post-treatment). α/β tubulin serves as the loading control. **e** Representative tracks from H3K27me3 CUT&Tag-sequencing for chromosome 1. **f** Representative tracks from H3K27me3 CUT&Tag-sequencing for chromosome 6. Tracks for control (siNeg) and H19 depleted (siH19) HCF cells are displayed in teal and yellow, respectively. The Refseq gene track is displayed in grey. Source data are provided as a Source Data file.
